# Key role of down-regulated in adenoma (*SLC26A3*) chloride/bicarbonate exchanger in linaclotide-stimulated intestinal bicarbonate secretion upon loss of CFTR function

**DOI:** 10.1101/2023.05.05.539132

**Authors:** Jessica B. Sarthi, Annie M. Trumbull, Shayda M. Abazari, Vincent van Unen, Joshua E. Chan, Yanfen Jiang, Jesse Gammons, Marc O. Anderson, Onur Cil, Calvin J. Kuo, Zachary M. Sellers

## Abstract

Duodenal bicarbonate secretion is critical to epithelial protection, nutrient digestion/absorption and is impaired in cystic fibrosis (CF). We examined if linaclotide, typically used to treat constipation, may also stimulate duodenal bicarbonate secretion. Bicarbonate secretion was measured in vivo and in vitro using mouse and human duodenum (biopsies and enteroids). Ion transporter localization was identified with confocal microscopy and de novo analysis of human duodenal single cell RNA sequencing (sc-RNAseq) datasets was performed. Linaclotide increased bicarbonate secretion in mouse and human duodenum in the absence of CFTR expression (*Cftr* knockout mice) or function (CFTR_inh_-172). NHE3 inhibition contributed to a portion of this response. Linaclotide-stimulated bicarbonate secretion was eliminated by down-regulated in adenoma (DRA, SLC26A3) inhibition during loss of CFTR activity. Sc-RNAseq identified that 70% of villus cells expressed *SLC26A3*, but not *CFTR*, mRNA. Loss of CFTR activity and linaclotide increased apical brush border expression of DRA in non-CF and CF differentiated enteroids. These data provide further insights into the action of linaclotide and how DRA may compensate for loss of CFTR in regulating luminal pH. Linaclotide may be a useful therapy for CF individuals with impaired bicarbonate secretion.

## INTRODUCTION

Proximal duodenal bicarbonate secretion, together with pancreatic bicarbonate secretion, is integral to preventing intestinal mucosal injury from gastric secretions and establishing a non-acidic intraluminal environment for activation of digestive enzymes. This importance is highlighted by cystic fibrosis (CF) intestinal disease and malnutrition, where decreased bicarbonate secretion is central to disease pathogenesis (1, 2). Duodenal bicarbonate secretion is highly regulated, driven by luminal acid and neurohormonal signaling, ultimately resulting in coordinated transepithelial bicarbonate secretion by the cystic fibrosis transmembrane conductance regulator (CFTR), chloride/bicarbonate exchange (e.g., down-regulated in adenoma, DRA; putative anion transporter-1, PAT-1), and/or decreased proton transport through sodium/hydrogen exchange (e.g. NHE3)(3). Despite its physiologic importance, there are limited therapies targeting duodenal bicarbonate secretion.

Linaclotide is a Food and Drug Administration (FDA)-approved medication for the treatment of constipation-type irritable bowel syndrome and chronic idiopathic constipation whose effect relies on its structural similarity to the heat-stable enterotoxin of *Escherichia coli* (STa). Both STa and linaclotide bind guanylyl cyclase C (GC-C) receptors on the apical surface of enterocytes and cause increases in CFTR-dependent chloride secretion and inhibition of NHE3-mediated sodium absorption (4–6). Previously, we identified that STa stimulates duodenal bicarbonate secretion through CFTR and CFTR-independent means (7, 8). Linaclotide has been investigated as a potential therapy for CF-related constipation (6, 9, 10), however, it is unclear if linaclotide stimulates duodenal bicarbonate secretion and if linaclotide could improve impaired intraluminal pH in CF patients by stimulating CFTR-independent bicarbonate secretion.

We undertook a series of experiments utilizing in vivo and in vitro bicarbonate secretion measurements in mice and humans to examine the effect of linaclotide on duodenal bicarbonate secretion. We coupled these experiments with scRNA-seq analysis of acid-based transporters in the human duodenum and confocal microscopy to characterize the brush border expression of the DRA chloride/bicarbonate exchanger in human duodenum in the presence or absence of CFTR activity and/or expression.

## RESULTS

### Linaclotide stimulates bicarbonate secretion in mouse and human duodenum

To determine linaclotide’s ability to stimulate intestinal bicarbonate secretion, we examined its effect on the duodenal segment of the small intestine, given the robust expression of acid-base transporters in this segment that function to neutralize the highly acidic gastric contents it encounters. We first examined the effect of in vivo intestinal perfusion of linaclotide on duodenal bicarbonate secretion. Linaclotide stimulated sustained increases in bicarbonate secretion at 10^-7^ M and 10^-5^ M (P<0.001 and P=0.004, n=9), with small and transient increases in bicarbonate secretion over baseline secretory rates at 10^-9^ M (P=0.033, n=9). There was no significant difference between 10^-7^ M or 10^-5^ M (P=0.926) (Figure 1A, B). These bicarbonate secretory rates were similar to those observed with STa (Figure 1B, 10^-9^ M: P=0.491 and 10^-7^ M: 0.792, respectively, n=11). Linaclotide did not stimulate net increases in luminal duodenal fluid secretion (10^-9^ M: 0.01 ± 0.05, 10^-7^ M: -0.05 ± 0.06, 10^-5^ M: -0.04 ± 0.05 ΔmL/cm•h, P=0.856, 0.445, 0.367, respectively, n=8), similar to what we have previously reported for STa in the duodenum (7). As linaclotide can decrease proton secretion by inhibiting NHE3 (6, 9), we examined if linaclotide actively stimulates bicarbonate secretion, or purely inhibits baseline proton secretion to raise intraluminal pH. Using the same in vivo technique, perfusion with S3226 (10^-5^ M), a selective inhibitor of NHE3, caused a significant increase in baseline bicarbonate secretion (3.78 ± 0.73 vs. 6.70 ± 0.68 μmol/cm•h, n=9-10, P=0.016). In the presence of S3226, linaclotide caused further increases in bicarbonate secretion (10^-9^ M: 2.22 ± 0.87, 10^-7^ M: 2.53 ± 0.82, 10^-5^ M: 2.38 ± 1.32 Δμmol /cm•h, n=10, P=0.075, P=0.032, P=0.237, respectively)(Figure 1C). In the presence of S3226, linaclotide-stimulated duodenal bicarbonate secretion was ∼1/3 of the response without S3226 (10^-7^ M: 37.9 ± 12.3%, 10^-5^ M: 35.6 ± 19.8%). Thus, in mouse duodenum, linaclotide alters duodenal luminal pH through a combination of inhibiting NHE3-mediating proton secretion and stimulating bicarbonate secretion.

**Figure 1.**
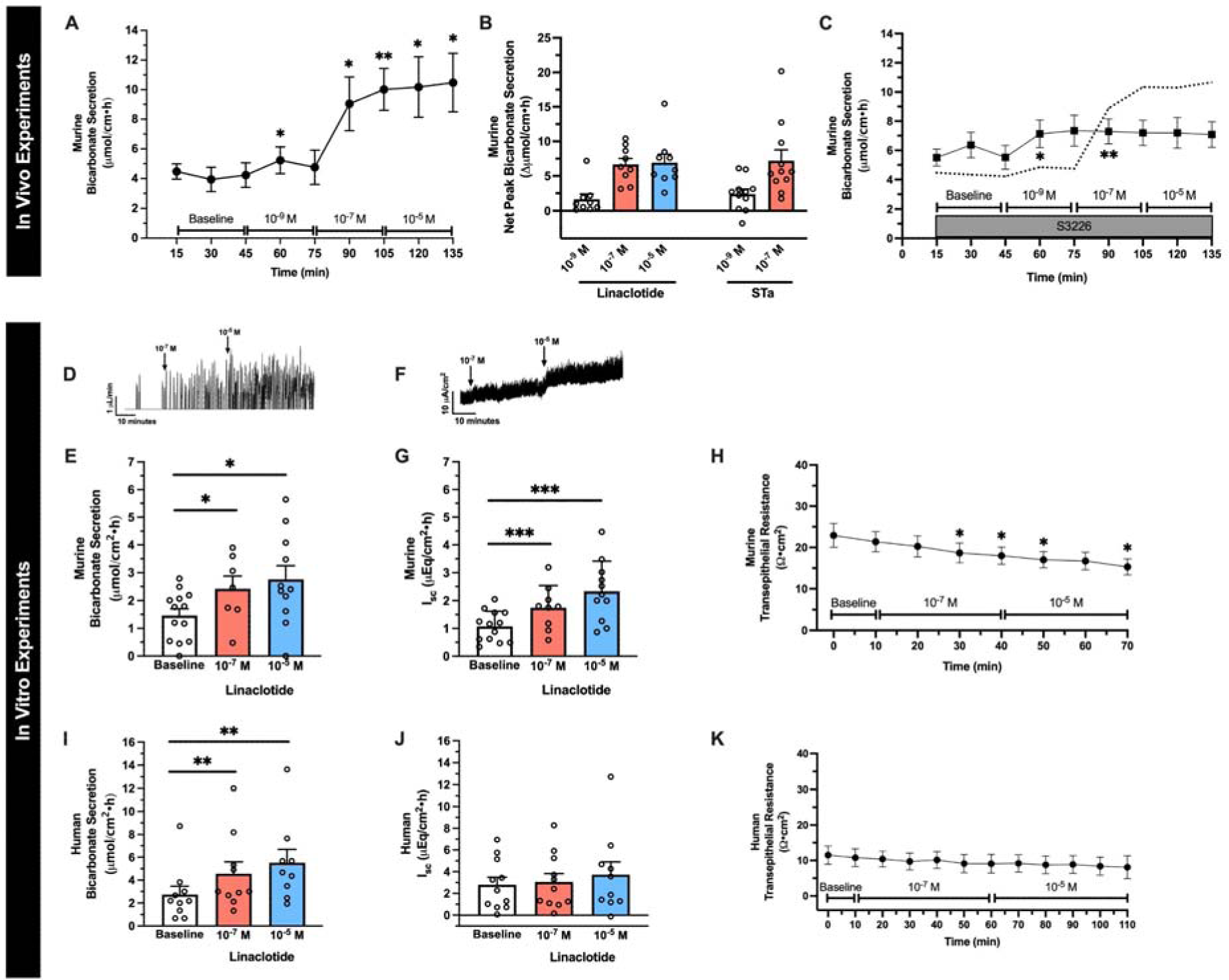
Linaclotide stimulates duodenal bicarbonate secretion in mice and humans. **A.** In vivo measurement of linaclotide-stimulated (10^-9^ M, 10^-7^ M, 10^-5^ M) duodenal bicarbonate secretion in C57Bl6/J mice (n=9). After 30-minute baseline measurements, each linaclotide dose was sequentially perfused for 30 minutes. *, P<0.05; **, P<0.01 vs. baseline by ANOVA. **B.** Net peak (max-baseline) bicarbonate secretion. Linaclotide values were from 1A. For STa (10^-9^ M, 10^-7^ M), separate experiments were performed (n=11). Each point represents a different mouse. **C.** In vivo effect of linaclotide (10^-9^ M, 10^-7^ M, 10^-5^ M) with NHE3 inhibitor S3226 (10^-5^ M) (n=10). Dotted line is the mean response to linaclotide (from 1A). *, P<0.05; **, P<0.01 vs. baseline by ANOVA. **D.** Representative trace for in vitro bicarbonate secretion in mouse duodenum. Y-axis is titrated HCl, which reflects base secreted by the tissue. During baseline there are few peaks. Upon stimulation, the number of titration events and/or volume of titrant increases. **E.** In vitro baseline and linaclotide-stimulated (10^-7^, 10^-5^ M, apical) mouse duodenal mucosal bicarbonate secretion. Each point (n=8-13) is a separate piece of duodenum from 10 mice. *, P<0.05 vs. baseline by ANOVA. **F.** Representative short-circuit current (I_sc_) trace for in vitro mouse experiments. **G.** In vitro baseline and linaclotide-stimulated (10^-7^, 10^-5^ M, apical) mouse duodenal I_sc_. Each point represents a separate piece of duodenum from the same mice in 1E. ***, P<0.001 vs. baseline by ANOVA. **H.** In vitro transepithelial resistance measurements from 1E and G. *, P<0.05 vs. baseline by ANOVA. **I-K.** Human duodenal mucosal bicarbonate secretion (I), I_sc_ (J), and transepithelial resistance (K) from endoscopic biopsies. Experiments were performed and data expressed similar as murine experiments in 1D-H. Each point (n=9-10) represents a different biopsy. **, P<0.01 vs. baseline by ANOVA. All data are means ± SEM, unless stated otherwise.

To further characterize linaclotide-stimulated bicarbonate secretion, we performed combined I_sc_ and bicarbonate secretory measurements in seromuscular-stripped mouse duodenum mounted in Ussing chambers. As seen in Figure 1D-G, this in vitro method showed similar results as the in vivo method. Linaclotide stimulated significant increases in duodenal bicarbonate secretion at 10^-7^ M (P=0.025, n=9) and 10^-5^ M (P=0.047, n=11), with no difference between them (P=0.875). Short-circuit current analysis revealed that linaclotide stimulated significant increases in I_sc_ at 10^-7^ M (P<0.001, n=9) and 10^-5^ M (P<0.001, n=11) (Figure 1F, G). Bicarbonate secretion and I_sc_ were accompanied by small, but significant (P<0.05) changes in transepithelial resistance (Figure 1H).

We next examined the potential for linaclotide to stimulate bicarbonate secretion in human duodenum using endoscopically obtained duodenal biopsies from subjects without known acid-base disturbances or histologic evidence of duodenal disease. Biopsies were mounted in Ussing chambers and combined bicarbonate secretion and I_sc_ measurements were performed, similar to Figure 1D-H. As seen in Figure 1I, 10^-7^ M and 10^-5^ M linaclotide stimulated significant increases in human duodenal bicarbonate secretion (P=0.006, n=11 and P=0.004, n=10, respectively), at similar magnitudes previously reported for STa in human duodenal biopsies (11). In contrast to mice, linaclotide did not stimulate a significant change in I_sc_ in human biopsies (10^-7^ M: P=0.071, n=11; 10^-5^ M: P=0.186, n=10) (Figure 1J) or transepithelial resistance (Figure 1J, K). Thus, linaclotide stimulates electrogenic bicarbonate secretion in mouse duodenum and electroneutral bicarbonate secretion in human duodenum.

### Linaclotide-stimulated duodenal bicarbonate secretion occurs independent of CFTR expression or function

Next, given our prior findings that STa can stimulate CFTR-independent duodenal bicarbonate secretion in mice (7), we studied linaclotide’s dependence on CFTR for bicarbonate secretion. We first repeated the in vivo bicarbonate secretion experiments by perfusing the duodenum with CFTR_inh_-172, a specific CFTR inhibitor.(12) Linaclotide continued to stimulate significant bicarbonate secretory responses at 10^-7^ M and 10^-5^ M (P=0.008 and P=0.018, respectively, n=12), but not 10^-9^ M (P=0.198, n=12), in the presence of CFTR_inh_-172 (2 × 10^-5^ M). In comparison, in vivo, CFTR_inh_-172 inhibited forskolin-stimulated (10^-4^ M) bicarbonate secretion by 65.1 ± 12.5% (P=0.022), preventing significant increases over baseline (P=0.778, n=6-9) (Supplementary Figure 1A and B). In vitro, sequential addition of CFTR inhibitors with different mechanisms of action, GlyH-101 (10^-5^ M, mucosal) and glibenclamide (3 × 10^-4^ M, mucosal), following CFTR_inh_-172 (2 × 10^-5^ M, serosal), resulted in no further increase in the inhibition of forskolin-stimulated I_sc_ (0.2% and -0.9 %, respectively). To expand upon these pharmacologic experiments, we performed in vitro Ussing chamber experiments with CFTR^tm1UNC^ knockout (KO) mice, which have global KO of the *Cftr* gene (13). Given the similar responses between 10^-7^ M and 10^-5^ M in prior experiments, for these studies we specifically focused on 10^-7^ M linaclotide. We found that linaclotide (10^-7^ M) stimulated significant increases in duodenal bicarbonate secretion in *Cftr* KO mice (P=0.028 vs. baseline, n=7). These magnitudes were similar to wildtype (WT) mice (P=0.728) (Figure 2B). In contrast to mice expressing CFTR, linaclotide stimulated minimal change in I_sc_ in *Cftr* KO mice (0.77 ± 0.31 ΔμA/cm^2^, P=0.049, n=8) with no change in transepithelial resistance (P=0.396) (Figure 2C, D), suggesting that in the absence of CFTR, linaclotide stimulates electroneutral bicarbonate secretion. To determine if these findings hold true in humans, we pre-treated human endoscopic duodenal biopsies with CFTR_inh_-172, which inhibits forskolin-stimulated I_sc_ in human duodenal enteroids (14)(Supplementary Figure 1C), and then performed repeat Ussing chamber measurements. Linaclotide (10^-7^ M) continued to stimulate significant increases in duodenal bicarbonate secretion in biopsies pre-treated with CFTR_inh_-172 (2 × 10^-5^ M, serosal, P=0.024, n=14), at a magnitude that was comparable to biopsies without CFTR_inh_-172 treatment (P=0.644) (Figure 2E). There was no difference in linaclotide-stimulated I_sc_ with or without CFTR_inh_-172 treatment (0.27 ± 0.11 vs. 0.13 ± 0.04 ΔμEq/cm^2^•h, P=0.239, n=11-14) (Figure 2F). Transepithelial resistance in the presence of CFTR_inh_-172 was unchanged before or after linaclotide stimulation (P=0.889, n=14) (Figure 2G).

**Figure 2.**
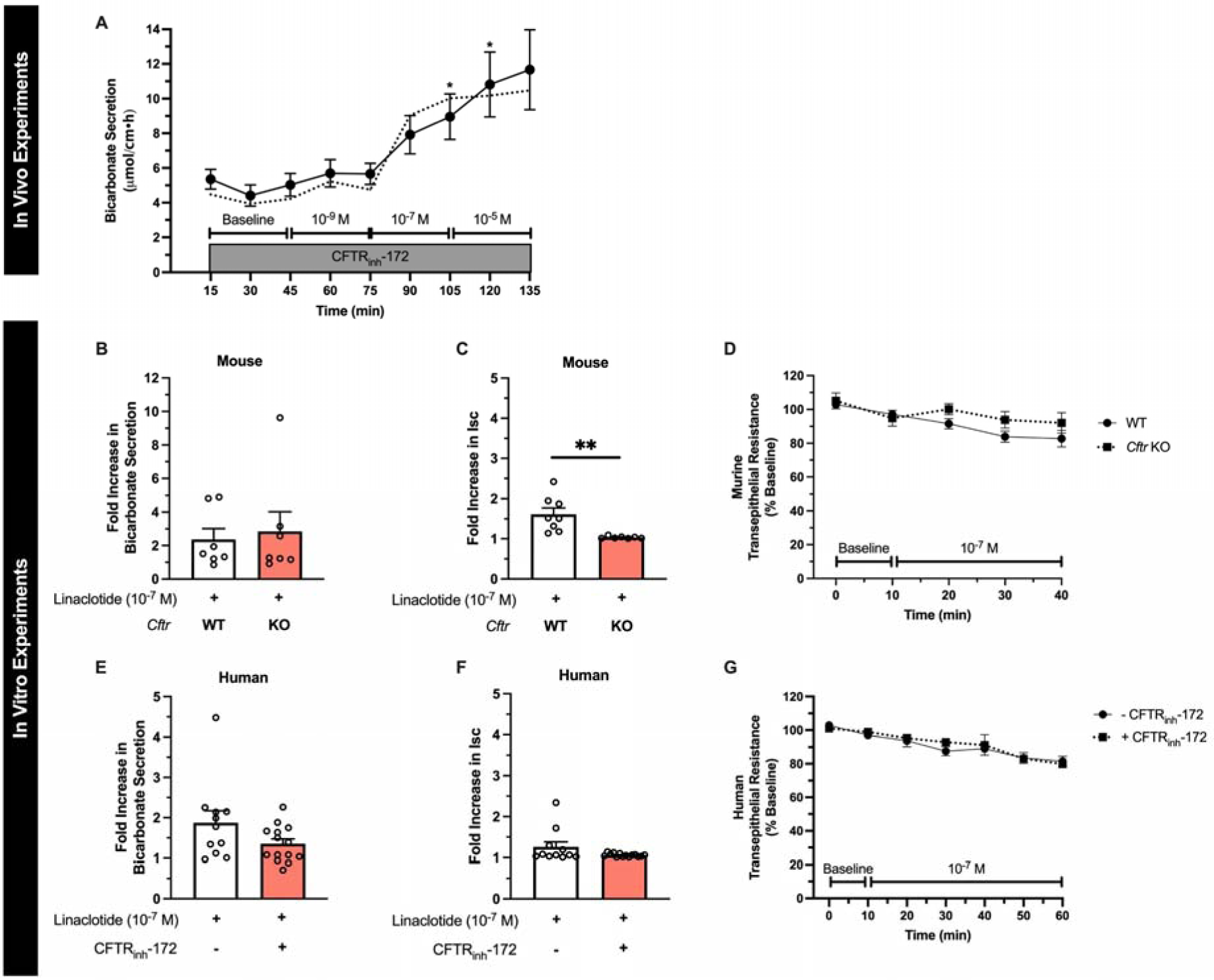
Linaclotide stimulates duodenal bicarbonate secretion independent of CFTR. **A.** In vivo measurement of duodenal bicarbonate secretion in mice, similar to Figure 1A, with the exception that each perfusate also contained CFTR_inh_-172 (2 × 10^-5^ M) (n=12). Dotted line is the mean response without CFTR_inh_-172 (from Figure 1A). *, P<0.05 vs. baseline by ANOVA. **B and C.** In vitro duodenal mucosal bicarbonate secretion (B) and I_sc_ (C) in wildtype and *Cftr* KO mice. Data are expressed as fold increase in linaclotide (10^-7^ M) stimulated responses over baseline bicarbonate secretion or I_sc_ in the same mouse. Each point (n=7) is a separate piece of duodenum from 5 mice. **, P<0.01 by unpaired Student’s t-test. **D.** Transepithelial resistance measurements in wildtype (WT) and *Cftr* KO mice. **E-G.** In vitro duodenal mucosal bicarbonate secretion (E), I_sc_ (F), and transepithelial resistance (G) in human endoscopic biopsies with or without CFTR_inh_-172 pre-treatment (2 × 10^-5^ M, 40-60 minutes). Each point (11–14) represents a different biopsy. All data are means ± SEM.

### Linaclotide-stimulated duodenal bicarbonate secretion upon loss of CFTR function is mediated by the DRA chloride/bicarbonate exchanger

To investigate the potential mechanism whereby linaclotide increases duodenal bicarbonate secretion independent of CFTR, we investigated the effect of DRA_inh_-A250 (selective inhibitor of DRA), DIDS (DRA-insensitive, non-selective anion exchanger inhibitor), and S3226 (selective inhibitor of NHE3) on duodenal bicarbonate secretion. Using our in vivo model, in separate experiments, each of these drugs were combined with CFTR_inh_-172 and perfused prior to stimulation with linaclotide. DRA_inh_-A250 (10^-5^ M) caused a 67 ± 8% reduction in CFTR-independent linaclotide-stimulated duodenal bicarbonate secretion at 10^-7^ M linaclotide (P=0.044, n=10), and 73 ± 6% inhibition at 10^-5^ M linaclotide (P=0.038, n=10) (Figure 3A-C). In contrast, DIDS (2 × 10^-4^ M) failed to significantly impact CFTR-independent linaclotide-stimulated duodenal bicarbonate secretion (10^-7^ M: -7 ± 14%, P=0.993, n=7; 10^-5^ M: -19 ± 14%, P=0.893, n=7). S3226 (10^-5^ M) reduced CFTR-independent linaclotide-stimulated duodenal bicarbonate secretion by 45 ± 19% (10^-7^ M) and 58 ± 16% (10^-5^ M); however, neither of these were statistically significant (P=0.254, P=0.136, respectively, n=9) (Figure 3A, B). Therefore, it appears that DRA is the primary source of bicarbonate transport for CFTR-independent linaclotide-stimulated duodenal bicarbonate secretion.

**Figure 3.**
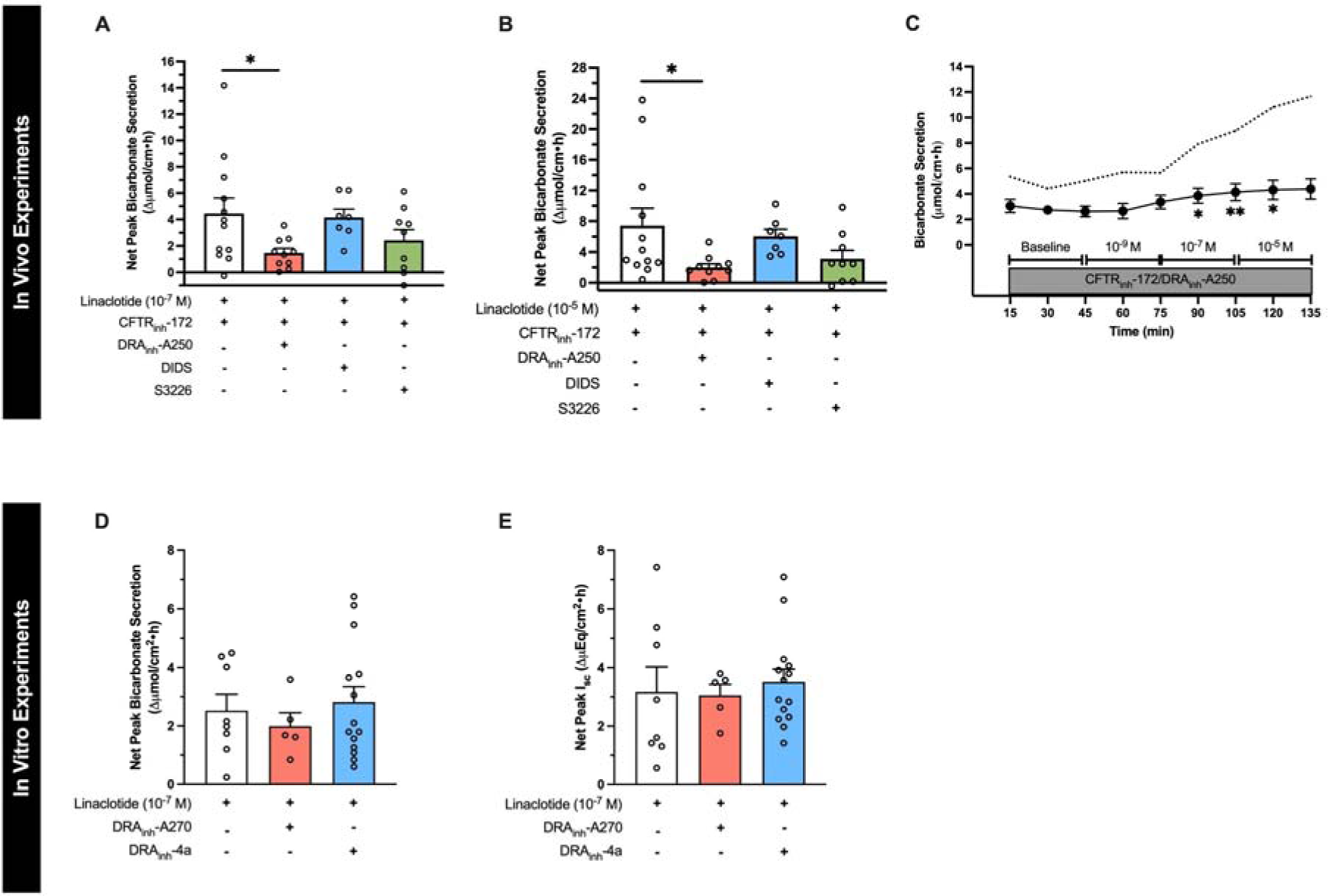
Critical role of DRA in CFTR-independent linaclotide-stimulated duodenal bicarbonate secretion. **A and B.** To determine the source of linaclotide-stimulated bicarbonate transport in the absence of CFTR function, in vivo experiments were repeated, similar to Figure 2A, except in addition to CFTR_inh_-172 (2 × 10^-5^ M, n=12), one of the following was also added to the luminal perfusate: DRA_inh_-A250 (10^-5^ M, n=10), DIDS (2 × 10^-4^ M, n=7), or S3226 (10^-5^ M, n=9). (A) linaclotide 10^-7^ M, (B) linaclotide 10^-5^ M. Each point represents a different mouse. *, P<0.05 vs. linaclotide + CFTR_inh_-172 (2 × 10^-5^ M) by ANOVA. **C.** Time course with linaclotide dose response (10^-9^ M, 10^-7^ M, 10^-5^ M) in the presence of CFTR_inh_-172 (2 × 10^-5^ M) and DRA_inh_-A250 (10^-5^ M), as indicated by circles and whiskers (n=12). Dotted line indicates mean response in the presence of CFTR_inh_-172 (2 × 10^-5^ M) only (from Figure 2A). *, P<0.05; **, P<0.01 vs. baseline by ANOVA. **D.** Net peak linaclotide-stimulated (10^-7^ M, apical) mouse duodenal mucosal bicarbonate secretion (D) and I_sc_ (E) from in vitro experiments, with or without DRA inhibition by DRA_inh_- A270 (10^-5^ M, bilateral, n=5) or DRA_inh_-4a (10^-5^ M, bilateral, n=14). Each point represents a separate piece of duodenum from 5-10 mice. All data are means ± SEM.

To determine if DRA’s critical role in linaclotide-stimulated duodenal bicarbonate secretion is unique to when there is functional loss of CFTR, we examined the role of DRA in mouse duodenal bicarbonate secretion measured in vitro. For these experiments, DRA_inh_-A270, a near identical chemical compound as DRA_inh_-A250 (Br^-^ to I^-^ substitution) with similar DRA specificity and inhibitory properties as DRA_inh_-A250 at 10^-5^ M (15), was used. DRA_inh_-A270 (10^-5^ M) had no significant inhibitory impact on linaclotide (10^-7^ M)-stimulated duodenal bicarbonate secretion (P=0.813, n=5-8) or I_sc_ (P=0.991, n=5-8) in the presence of CFTR function (Figure 3D and E). In vivo testing also showed no inhibitory effect of DRA_inh_-A270 on linaclotide-stimulated bicarbonate secretion without CFTR inhibition (DMSO + linaclotide 10^-7^ M: 4.3 ± 1.1 vs. DRA_inh_- A270 + linaclotide 10^-7^ M: 3.5 ± 0.3 μmol.cm^-1^.h^-1^, n=4 each, P=0.542). To confirm lack of effect of DRA inhibition under these conditions, we performed additional in vitro experiments using DRA_inh_-4a, a DRA specific inhibitor that is structurally and mechanistically distinct from DRA_inh_- A250 and DRA_inh_-A270 (16). These experiments showed similar results as DRA_inh_-A270; no inhibition of linaclotide-stimulated duodenal bicarbonate secretion (P=0.899, n=8-16) or I_sc_ (P=0.870, n=8-16) in the presence of CFTR function. Thus, CFTR is sufficient for linaclotide-stimulated bicarbonate secretion, but upon loss of CFTR function, DRA appears to compensate for its loss.

### DRA expression in human duodenum

DRA and PAT-1 are the primary chloride/bicarbonate exchangers expressed in mouse intestine, with PAT-1 showing dominant RNA expression in the small intestine and DRA RNA being more abundant in the colon.(17) There is limited data regarding DRA expression in human duodenum.(18, 19) We capitalized on two previously published human duodenum scRNA-seq datasets(20, 21) to examine DRA mRNA expression in human duodenum. The Elmentaite et al. dataset(21) contained 5,944 cells with an average of 4,164 reads per cell from 5 healthy adult duodenal samples, allowing for high confidence in cell-type separation. The Busslinger et al. dataset(20) contained 702 crypt cells and 923 villus cells from 2 adult duodenal samples, with an average of 10,381 reads per cell. Data from one crypt sample (“Crypt 1”) was excluded due to very low sequence reads (2,088 per cell). We verified accurate classification into crypt and villus cells by examining *LGR5* expression and proliferation markers *MKI67* and *PCNA*, which are generally restricted to the crypt region. These were all more abundant in the crypt group, with *LGR5* being exclusively expressed in the crypt, but not villus, group (Supplementary Figure 2A-C). Using these two datasets, we examined expression of the key acid-base transporters in the duodenum, DRA (*SLC26A3*), PAT-1 (*SLC26A6*), *CFTR*, and NHE3 (*SLC9A3*). Our analysis showed that *SLC26A3* mRNA is predominantly expressed in enterocytes (Figure. 4A), in both the crypt and villus groups (Figure 4E), with higher expression and expressed in more cells than *SLC26A6*, *CFTR*, or *SLC9A3* (Figures 4A-D and E-H). Of note, Crypt 2 had lower *SLC26A3* expression than Crypt 3, which correlated with higher *CFTR, MKI67, PCNA* expression, more *LGR5*+ cells, and less *SLC9A3*+ cells, suggesting Crypt 2 contained more cells deeper within the crypts than Crypt 3 (Figures 4E, G, H and Supplementary Figure 2). We confirmed DRA protein expression in human duodenal biopsies (n=3) using immunofluorescence imaging. The majority of DRA is localized to the intracellular apical region, with a smaller fraction present at the apical brush border (Figures 4I-K).

**Figure 4.**
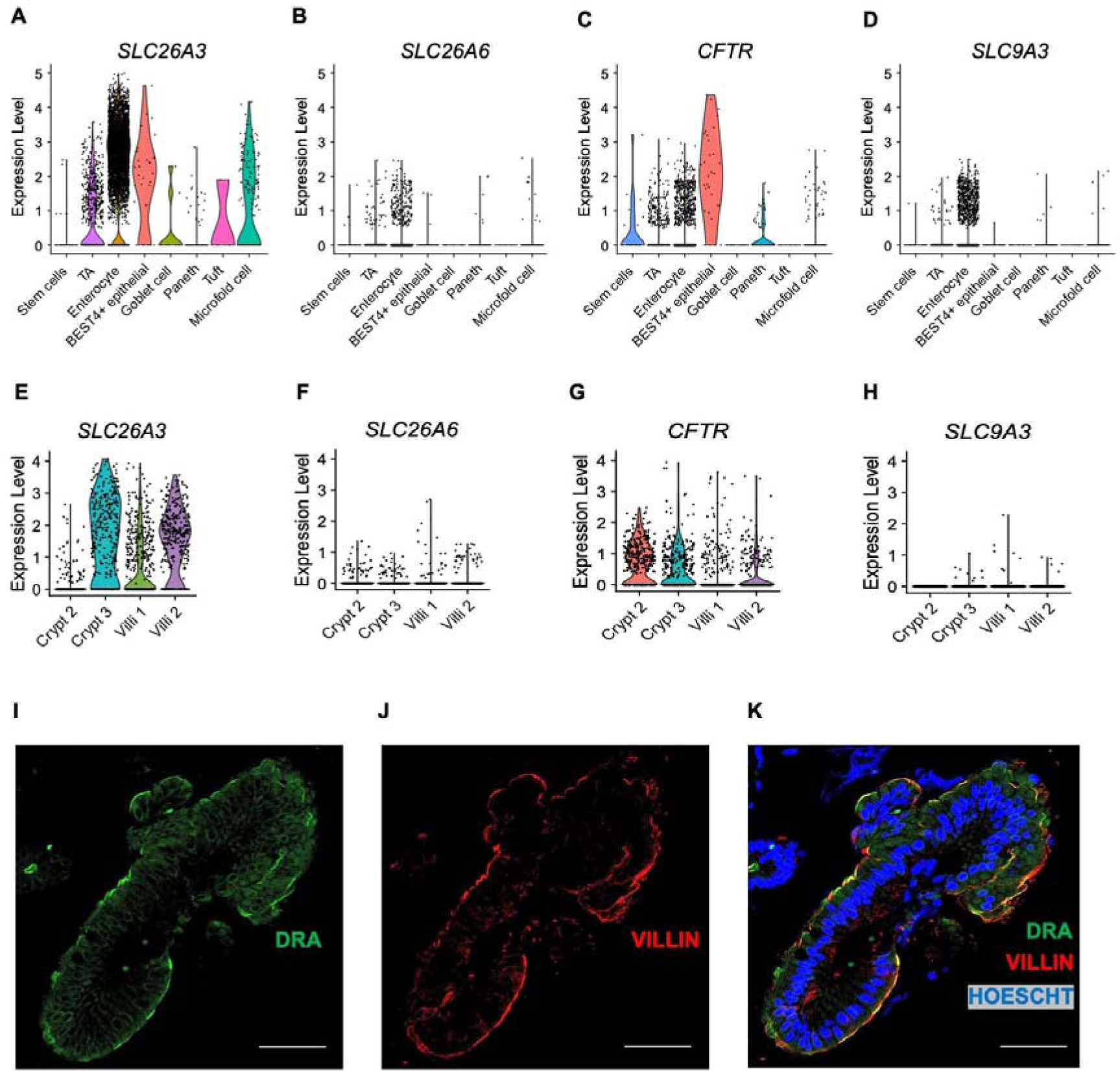
Cellular and membrane expression of *SLC26A3* (DRA) in human duodenal enterocytes. **A-H.** Cellular mRNA expression of *SLC26A3* (DRA), *SLC26A6* (PAT-1), *CFTR*, and *SLC9A3* (NHE3) based on re-analysis of Elmentaite et al.(21) (A-D) and Busslinger et al.(20) (E-H). Violin plots represent expression relative to all cells of that type, with each point representing the expression of individual cells within each type. **I-K.** Representative confocal immunofluorescence imaging of DRA (I), villin (marker of apical brush border, J), and nucleus (K) in human duodenum from endoscopic biopsy (n=3). Scale bar = 20 μm.

Given DRA’s functional independence from CFTR in mediating linaclotide-stimulated duodenal bicarbonate secretion, we quantified the proportion of *SLC26A3*-expressing cells that co-expressed *CFTR*. In the Elmentaite et al. dataset, 88.8% of *SLC26A3*-expressing enterocytes co-expressed *SLC26A3* and *CFTR* (Figure 5a). Using the Busslinger et al. dataset to examine crypt vs. villus localization, 63.0% of *SLC26A3* positive crypt cells co-expressed *SLC26A3* and *CFTR*, whereas in the villus cells, only 29.6% did so (Figures 5B and C). These findings were not ubiquitous for all chloride/bicarbonate exchangers. *SLC26A6* and *CFTR* co-expression was relatively low in enterocytes (16.7%) (Supplementary Figure 3A), but examination of the crypt and villus cell types showed a higher percentage of *CFTR* co-expression with *SLC26A6* than *SLC26A3*, with 81.8% of crypt cells and 42.9% of villus cells co-expressing *SLC26A6* and *CFTR* (Supplementary Figures 3B and C). In the Elmentaite et al. dataset, *SLC9A3* and *CFTR* were co-expressed in 14.3% of enterocytes, similar to *SLC26A6* (Supplementary Figure 3D). The Busslinger et al. dataset possessed insufficient numbers of *SLC9A3*-expressing cells to analyze *SLC9A3* and *CFTR* co-expression in crypts and villi.

**Figure 5.**
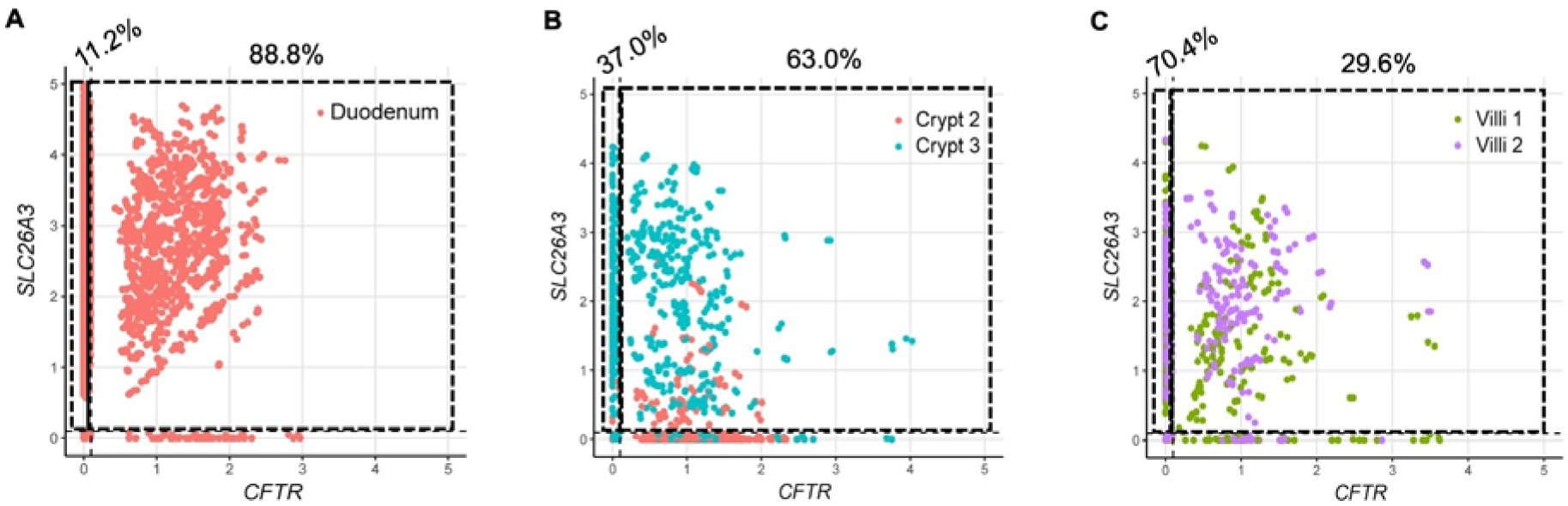
*SLC26A3* and *CFTR* co-expression in human duodenum by single cell RNA sequencing. **A-C.** Co-expression of *SLC26A3* (DRA) and *CFTR* mRNA using FeatureScatter based on Elmentaite et al.(21) enterocytes (A) and Busslinger et al.(20) crypt and villi (B and C) datasets that were analyzed in Figure 4. Numbers on top of graphs represent the percentage of *SLC26A3*-expressing cells that express *SLC26A3* only (left) or *SLC26A3* and *CFTR* (right).

### Linaclotide alters the apical localization of DRA

Ion channel activity can be regulated by altering membrane expression of channels. We examined if linaclotide alters DRA activity by changing its apical brush border expression. For these experiments we utilized human derived three-dimensional duodenal enteroids. Given that the linaclotide receptor GC-C is expressed at the apical surface, we generated reverse polarity apical-out duodenal enteroids, which allow facile access to the apical membrane via the enteroid exterior (Supplementary Figures 4A and B).(22) Using qPCR, we verified that apical-out enteroids express *CFTR*, *SLC26A3, SLC26A6, SLC9A3, and GUCY2C* in undifferentiated and 3-day differentiated enteroids (Supplementary Figures 4C and D. 4). As expected, *LGR5* was decreased in differentiated apical-out enteroids. Using this model of human duodenum, we incubated apical-out differentiated enteroids with linaclotide (10^-7^ M, 40 minutes) or vehicle control (water, 40 minutes) and performed confocal immunofluorescence to localize DRA. As seen in Figures 6A-D, linaclotide significantly increased apical brush border expression of DRA (n=12-16). In contrast, when we performed the same experiment with NHE3 localization, we found no change in the apical brush border expression of NHE3 upon linaclotide stimulation (Supplementary Figures 5A-D). We verified that myosin VI (*MYO6*), a key protein in NHE3 trafficking,(23) mRNA was present in apical-out enteroids. *MYO6* mRNA was expressed at similar levels in this model as many of the other transporters we examined (Supplementary Figures 4C and D). To determine if linaclotide also stimulates DRA localization upon lack of functional CFTR, we repeated the experiments in the presence of CFTR_inh_-172 (2 × 10^-5^ M, 40 minutes). Compared to vehicle control (DMSO, 40 minutes), CFTR inhibition resulted in an increase in the apical expression of DRA (P=0.032, n=26-31)(Figures 6E-H), but not NHE3 (P=0.633, n=16-30) (Supplementary Figures 5E-H). Of note, DMSO alone increased DRA brush border expression compared to water (MFI means ± SEM: 10.94 ± 1.32 vs. 3.43 ± 1.03, n=11-15, P<0.001). Linaclotide stimulation in the presence of CFTR_inh_-172 did not alter DRA or NHE3 brush border expression (P= 0.730, n=31; P=0.535, n=29-30, respectively) (Figures 6E-G and I, Supplementary Figures 5E-G and I).

**Figure 6.**
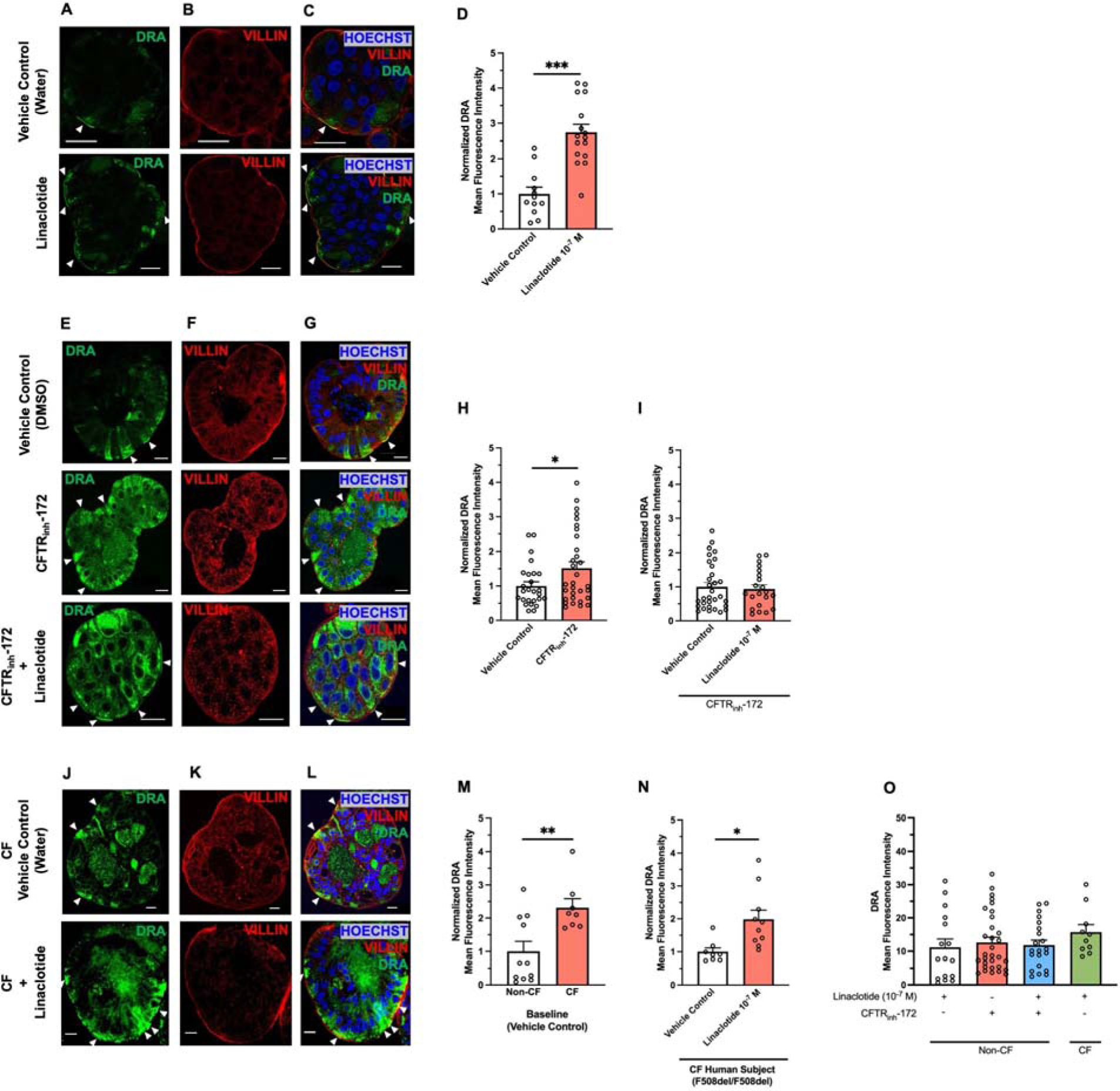
Linaclotide increases membrane expression of DRA in apical-out human duodenal enteroids. **A.** Representative confocal microscopy images of DRA (A), villin (B), and DRA and villin and Hoescht (C) during control conditions (water, 40 minutes, top) or linaclotide (10^-7^ M, 40 minutes, bottom). **D-F.** Quantification of DRA present at the apical brush border using villin to define this cell region. Apical brush border DRA mean fluorescence intensity following linaclotide (10^-7^ M, 40 minutes, n=16 ) treatment was normalized to vehicle controls for comparison.CFTR_inh_- 172 (2 × 10^-5^ M, 40 minutes) and/or linaclotide (10^-7^ M, 40 minutes) treatment was normalized to vehicle controls for comparison. Vehicle control for linaclotide was water and for CFTR_inh_-172 was DMSO. Columns with whiskers are mean ± SEM with each dot representing a different enteroid. Enteroids from three different patients were used for each condition. Significance determined by unpaired Student’s t-test. For enteroids treated with both CFTR_inh_-172 and linaclotide, enteroids were pretreated with CFTR_inh_-172 for 40 minutes prior to linaclotide treatment. **G and H.** DRA membrane expression in non-CF and CF (F508del homozygous) enteroids during vehicle control conditions (G and H, water) or during linaclotide stimulation (H, 10^-7^ M, 40 minutes). N=8-11 enteroids from 1 patient across 2 different passages. Data were normalized to non-CF enteroids (G) or vehicle control in CF enteroids (H). Significance determined by unpaired Student’s t-test. **I.** To compare DRA membrane expression across non-CF and CF enteroids in different situations, absolute DRA membrane mean fluorescence intensity was plotted. Data are from experiments performed in D-H. Significance determined by ANOVA. Arrows provided to assist with identifying regions of interest. All data are means ± SEM, unless stated otherwise.

To determine how our CFTR_inh_-172 studies compare to chronic loss of CFTR expression and function, we examined DRA brush border expression in human duodenal enteroids isolated from a CF subject with F508del homozygous *CFTR* variants. Comparison of baseline (water vehicle control, 40 minutes) DRA brush border expression between non-CF and CF enteroids showed CF enteroids had increased DRA expression at the apical brush border (Figures 6J-M, n=8-11, P=0.006). Linaclotide stimulation (10^-7^ M, 40 minutes) increased DRA brush border expression in F508del homozygous CF enteroids, compared to CF enteroids treated with vehicle control (water, 40 minutes)(Figures 6J-L and N, n=8-10, P=0.010). To examine the discrepancies between CFTR_inh_-172 + linaclotide in non-CF enteroids and linaclotide in the CF enteroids, we examined the absolute mean fluorescent intensities (MFI) of each condition. We found that CFTR_inh_-172 alone caused similar increases in absolute DRA brush border expression as linaclotide in non-CF enteroids (with or without CFTR_inh_-172) or CF enteroids (Figure 6O, n=10-31, P=0.676), suggesting that lack of additional DRA membrane expression in non-CF enteroids treated with CFTR_inh_-172 + linaclotide may be due to maximal DRA brush border expression already being achieved upon CFTR_inh_-172 treatment. Taken together with our bicarbonate secretory measurements, these data show that linaclotide can alter DRA activity and/or brush border expression, in the presence or absence of functional CFTR. In contrast, linaclotide appears to modulate NHE3 activity, rather than brush border expression, regardless of CFTR function.

### Inhibition of CFTR transiently increases intracellular pH

To investigate why CFTR inhibition increases DRA brush border expression, we examined the effect of CFTR_inh_-172 on duodenal enteroid intracellular pH (pH_i_). We hypothesized that loss of CFTR activity would lead to intracellular bicarbonate trapping (resulting in increased pH_i_) and DRA may traffic to the apical brush border as part of the cell’s effort to restore bicarbonate transport and normalize the altered pH_i_. Undifferentiated and 3-day differentiated enteroids grown on coverslips and CFTR_inh_-172-induced (or DMSO) changes in pH_i_ were monitored with live cell fluorescence imaging. CFTR_inh_-172 (2 × 10^-5^ M) resulted in significant increases in pH_i_ in undifferentiated and differentiated enteroids, although the slopes and magnitudes of these increases were different (Figure 7, n=57-89, P<0.001), with undifferentiated enteroids showing more rapid and greater pH_i_ changes (P<0.001). DMSO (1:1000) had no effect on pH_i_ (data not shown). Following CFTR_inh_-172, all enteroids underwent pH_i_ recovery. Pre-treatment with DRA_inh_-A250 (10^-5^ M, 5 min) did not alter the CFTR_inh_-172-induced increase in pH_i_ but did alter the pH_i_ recovery. In undifferentiated enteroids, DRA_inh_-A250 decreased the slope and magnitude of pH_i_ recovery (Figures 7A-C, n=31, P<0.001). In differentiated enteroids, the magnitude of pH_i_ recovery was not different with DRA_inh_-A250 (Figure 7F, n=37, P=0.213), however, the recovery slope was altered, albeit in a different way than undifferentiated enteroids (Figure 7D and E). Thus, inhibition of CFTR function causes an increase in pH_i_, the recovery of which is at least in part influenced by DRA activity.

**Figure 7.**
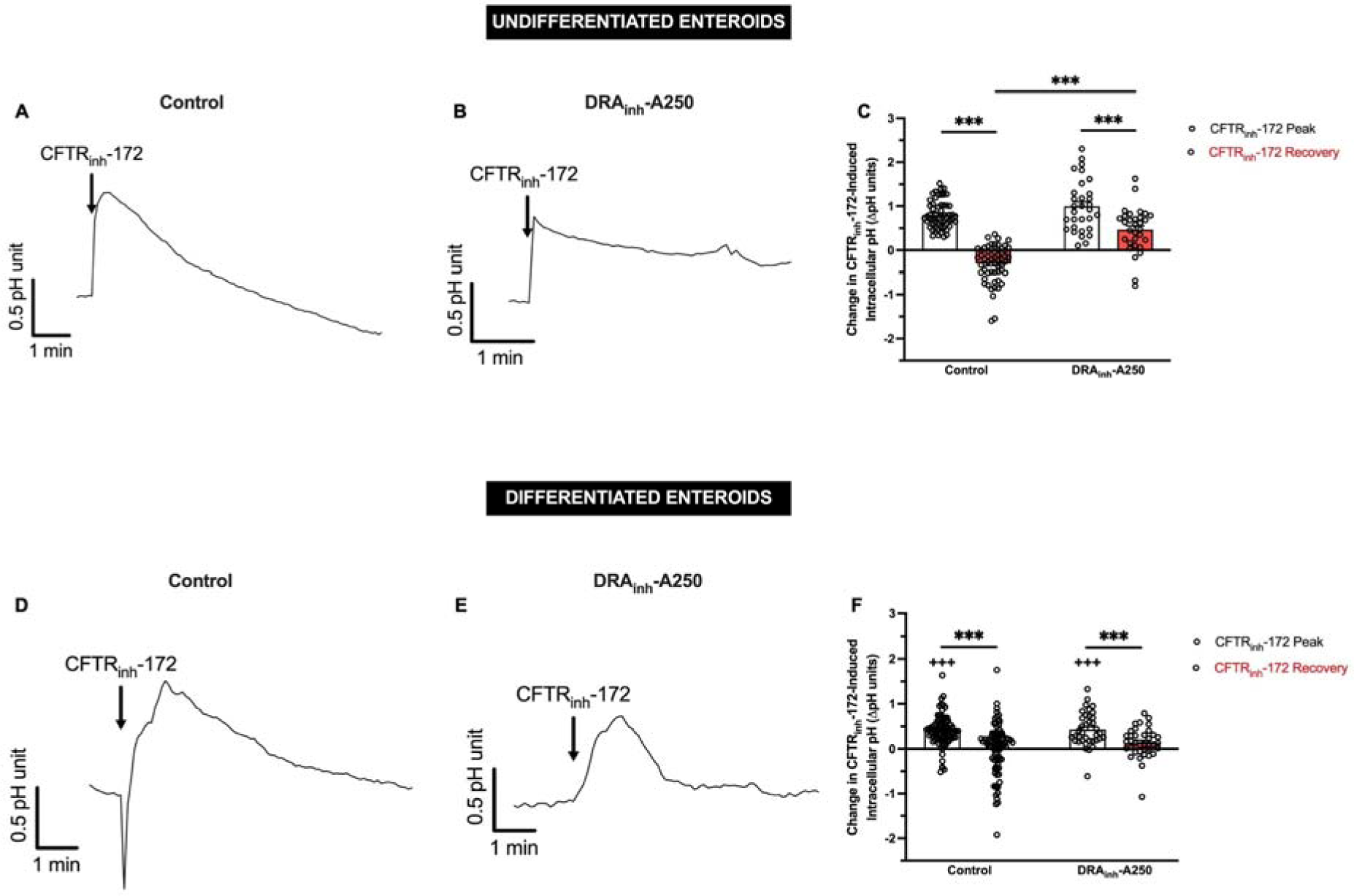
Inhibition of CFTR activity induces increases in intracellular pH. **A and B.** Mean timecourse of CFTR_inh_-172 (2 × 10^-5^ M) induced changes in pH_i_ in the absence (A) or presence (B) of DRA_inh_-A250 (10^-5^ M) pre-treatment in undifferentiated human duodenal enteroids. **C.** Quantitative comparison of the peak change (peak – baseline) or recovery (plateau – baseline) in pH_i_ units for enteroids exposed to CFTR_inh_-172 alone or DRA_inh_-A250 then CFTR_inh_-172. Data are expressed as means ± SEM**. D-F.** Similar experiments and analyses were performed in 3-day differentiated enteroids as A-C. Each circle represents a different cell within an enteroid. Measurements were taken from multiple enteroids across multiple coverslips. ***, P<0.001 by paired (within control or DRA_inh_-A250 groups) or unpaired (across control or DRA_inh_-A250 groups) Student’s t-test. +++, P<0.001 compared to undifferentiated enteroids by unpaired Student’s t-test.

## DISCUSSION

Numerous studies in animal models and humans have shown the critical importance of duodenal bicarbonate secretion in maintaining epithelial integrity and creating the appropriate intestinal intraluminal pH for digestion. In CF, with genetic loss of *CFTR*, or *Helicobacter pylori* infection, with down-regulation of CFTR and PAT-1,(24) decreased duodenal mucosal bicarbonate secretion contributes to malabsorption and duodenal ulcers, respectively. While eradication of *Helicobacter pylori* with antibiotics may restore bicarbonate function, the most common way to improve small intestinal pH in CF is through chronic gastric acid suppression. Ivacaftor, a CFTR potentiator, was shown to improve proximal intestinal pH in CF patients with the G551D CFTR gating mutation (accounts for 4-5% of *CFTR* disease-causing variants).(25) While additional CFTR modulators are FDA-approved, it remains unknown whether these positively impact small intestinal pH. This gap, coupled with lack of potential bicarbonate-correcting therapy for CF patients ineligible for CFTR modulator therapies (due to genotype or medical contraindication), has driven us to seek ways to improve intestinal bicarbonate secretion independent of CFTR. Our current data are in line with prior mouse and human duodenum studies with STa (7, 11) and a recent study by Tan et el. showing that linaclotide can stimulate jejunal bicarbonate secretion in mice.(9) Uniquely, we have shown that linaclotide stimulates both mouse and human bicarbonate secretion in the duodenum, where bicarbonate secretion is physiologically important. Furthermore, we have provided evidence for the roles (or lack of) membrane trafficking of DRA and NHE3 in this process and how decreases in CFTR function may impact cellular pH.

Linaclotide represents an attractive target for restoration of deficient duodenal bicarbonate secretion in CF. Our data in both mice and humans provide greater confidence in the translatability of our results to clinical care. While we did not study other segments of the small intestine, linaclotide stimulates jejunal bicarbonate secretion in F508del CF mice.(9) Braga Emidio et al. identified that linaclotide’s half-life in small intestinal fluid was 48 minutes,(26) suggesting that its effect is likely to be most influential in the proximal small intestine. Comparing linaclotide-stimulated and acid-stimulated duodenal bicarbonate secretion using the same methodology,(27, 28) the linaclotide effect is about 75-100% of the stimulatory effect of acid, indicating linaclotide could produce a clinically meaningful impact for patients. With existing FDA approval, and the use of clinically relevant dosing,(9) our results may provide immediate application to patients.

In the absence of CFTR, the logical targets for intestinal pH modulation are either 1) increasing chloride/bicarbonate exchanger activity, and/or 2) decreasing sodium/hydrogen exchange. Linaclotide increases intraluminal small intestinal fluid in mice by inhibiting NHE3-dependent sodium absorption.(6, 9) Another byproduct of NHE3 inhibition is a decrease in proton secretion, which can raise intraluminal pH. Consistent with this, NHE3 inhibition in our studies “increased bicarbonate secretion,” likely by producing less luminal protons to neutralize secreted bicarbonate anions. In our NHE3 trafficking studies, we did not see a significant decrease in NHE3 membrane localization with linaclotide treatment. NHE3 activity may be impacted by shifting along the microvilli,(23) however, our imaging resolution was unable to determine these potential nanometer shifts. Nonetheless, our in vivo data supports that NHE3 inhibition alone does not account for linaclotide’s ability to increase duodenal pH. Our studies using specific DRA inhibitors and DRA protein trafficking studies in apical-out human enteroids, indicate that DRA-mediated chloride/bicarbonate exchange is critical to linaclotide stimulated duodenal bicarbonate secretion upon loss of CFTR activity. This helps explain the lack of linaclotide-stimulated short-circuit current in *Cftr* KO mice, human biopsies with CFTR_inh-_172, and prior studies by McHugh et al. in F508del mice.(6) Interestingly, we found that CFTR inhibition alone caused an increase in DRA membrane trafficking. There was also increased apical membrane DRA expression in F508del homozygous duodenal enteroids even without stimulation. In mouse intestinal crypts, Strubberg et al. identified that pH_i_ was raised in *cftr* KO mice or wildtype mice treated with CFTR_inh_-172.(29) In fact, ∼20 years ago, Kaunitz et al. proposed the “CF Paradox” whereby diminished bicarbonate transport may increase intracellular buffering in CF and prevent duodenal ulcer formation despite diminished bicarbonate secretion.(30) Thus, we hypothesized that upon CFTR blockade (or genetic lack of *CFTR*), the rise in pH_i_ may trigger an increase in DRA expression and protein localization to at least partially restore bicarbonate secretion and limit alterations in pH_i_. Our current pH_i_ data with CFTR_inh_-172, supports this hypothesis. Bijvelds et al. reported a nearly 2,000-fold increase in *DRA* mRNA in a CF ileal biopsy compared to non-CF ileum (n=1 each).(31) The lack of additional membrane trafficking of DRA by linaclotide in the presence of CFTR inhibition may indicate that DRA trafficking requires CFTR function or that a ceiling affect for DRA membrane expression has already been achieved. Based on our current data we favor the latter interpretation since linaclotide increased DRA trafficking in F508del homozygous CF enteroids and a comparison of MFIs across conditions showed that while the relative magnitude of DRA trafficking may differ between vehicle controls/conditions, the absolute magnitudes are similar. While CFTR inhibition caused similar pH_i_ changes in undifferentiated (crypt-like) and differentiated (villus-like) enteroids, how DRA inhibition impacted pH_i_ recovery was different. This may be due to increased expression of a larger variety of acid-base transporters in differentiated enteroids that may also contribute to pH_i_ normalization.(19) Further study on how the expression and activities of DRA and other acid-base transporters are altered in CF intestine, especially in human samples, may identify new potential therapies for CF intestinal disease.

Prior to our study, our knowledge on the intestinal expression profile of DRA and other acid-base transporters was largely based on mouse studies. Wang et al. reported high expression of *Slc26a3* (DRA) RNA in the colon, but negligible RNA expression in the small intestine (duodenum, jejunum, ileum). Conversely, *Slc26a6* (PAT-1) RNA expression was high in mouse duodenum, jejunum, and ileum, with minimal in the colon.(17) Yet, *Slc26a3* KO mouse studies suggest a prominent role for DRA in basal and stimulated bicarbonate transport. (32, 33) Human studies are sparse. Yin et al. used human duodenal enteroids to examine *SLC26A3*, *SLC26A6*, and *SLC9A3* (NHE3) mRNA expression in undifferentiated and terminally differentiated human duodenal enteroids.(19) Taking advantage of published scRNA-seq datasets from the human duodenum, we have provided a comparison on the expression of *SLC26A3*, *SLC26A6*, and *SLC9A3*, from human duodenum tissue. *SLC26A3* mRNA was generally expressed ∼3-4x higher than *SLC26A6*, *CFTR*, and *SLC9A3*. In the *CFTR* high expressing *BEST4*+ cells, *CFTR* expression levels approached that of *SLC26A3*. Of note, *SLC26A3* was also highly expressed in *BEST4*+ cells, although enterocytes comprise the majority of *SLC26A3*-expressing cells. These findings, put in context of prior studies, suggest a more prominent role for DRA in the human duodenum than may have generally been appreciated from mouse studies alone. Future work examining not only the mRNA expression, but also protein expression, and especially pertinent to membrane ion transporters, membrane localization of DRA, PAT-1, and NHE3 across different segments of human intestine may advance our knowledge regarding the regional roles of these acid-base transporters in health and disease.

A potential limitation of our study is the general reliance on pharmacologic inhibitors throughout our experiments. In *Xenopus levis* oocytes, CFTR_inh_-172 (2 × 10^-5^ M) inhibited ∼70% of human CFTR current in patch-clamp experiments (34). In our experiments, CFTR_inh_- 172 (2 × 10^-5^ M) significantly inhibited CFTR ion transport in mouse tissue and human enteroids, with observed results similar to that by Yin et al.(19) Complementary results in *Cftr* KO mice and CF enteroids provide further confidence in our findings with CFTR_inh_-172. For DRA, we did not perform experiments in *Slc26a3* KO animals or cells, however, we did use 3 different DRA inhibitors, from 2 different structural categories and modes of inhibition. While confidence in pharmacological inhibitors can wane with time and increased use, we believe the consistency of results across these different inhibitors provides confidence in the interpretation of the data.

In summary, we have identified that linaclotide may have more pleiotropic ion transport effects beyond CFTR- and NHE3-mediated chloride and sodium transport, respectively. We have shown, using mice, duodenal biopsies, and human duodenal enteroids, that linaclotide stimulates bicarbonate secretion, and does so at least in part, through DRA recruitment to the apical membrane. Our findings that this remains possible independent of CFTR activity and/or expression has potentially important implications for CF patients, who may benefit from linaclotide’s bicarbonate stimulatory effect, in addition to its inhibitory effect on NHE3, even if they can’t leverage its pro-chloride secretory properties.(6) Furthermore, our identification of high DRA expression in the human duodenum may spur interest in identifying DRA-activating drugs targeting proximal small intestinal disease.

## METHODS

### Chemicals

CFTR_inh_-172 was purchased from Cayman Chemical (Ann Arbor, MI), 4,4’- Diisothiocyanatostilbene-2,2’-disulfonate (DIDS) and S3226 were purchased from Sigma-Aldrich (St. Louis, MO). DRA inhibitors (DRA_inh_-A250, DRA_inh_-A270, DRA_inh_-4a) were synthesized and purified as previously described.(15, 16, 35) Linaclotide was purchased from Santa Cruz Biotechnology, Inc (Dallas, TX).

### Animals

Adult (8-16 week-old) C57BL/6J mice from Jackson Laboratories (Bar Harbor, ME) were used as wildtype control mice. CFTR^tm1Unc^ knockout (KO) mice were obtained from Case Western University CF Mouse Models Core Facility and maintained on 50% PEG-3350 with electrolytes solution in their drinking water with standard chow. Similar number of male and female mice were used in experiments.

### In Vivo Measurement of Bicarbonate Secretion

In vivo measurement of duodenal bicarbonate secretion was performed using a well-validated technique.(36) Animals were maintained with free access to food and water for up to 1 hour before the experiment, then fasted from chow. Mice were anesthetized with oxygen-delivered isoflurane (1-3%) at 1 L/minute via a vaporizer (Braintree Scientific, Inc, Braintree, Mass). Mouse temperature was monitored by rectal probe and maintained at 37°C through automated warming using a controlled warming pad (ATCC 2000, World Precision Instruments, Sarasota, FL). Anesthetic plane was assessed by respiratory rate and toe pinch reflex. Respiratory and heart rates were determined every 15 minutes. Animals could be sustained for more than 3 hours under these experimental conditions. To isolate the proximal duodenum, the abdomen was opened by a central vertical incision, and the proximal 5–10 mm of duodenum, from the pylorus to just proximal to the entry of the common bile duct, was isolated in situ without compromising vascular supply. A small polyethylene tube (PE-50) with a distal flange was advanced to the duodenal bulb via the stomach, and a ligature was secured around the pylorus. A distal intestinal incision was made, and PE-50 flanged tubing was advanced to just proximal to the entry of the pancreaticobiliary duct to prevent entry of the pancreatic or biliary secretions and allow for collection of effluent from isolated segment. The isolated duodenal segment was gently flushed and then continuously perfused (Harvard Infusion Pump, Harvard Apparatus, South Natick, MA, USA) at a rate of 0.21 mL/min with 154 mM NaCl (37°C) ± stimulatory/inhibitor drugs. Effluents from the isolated segment were visually free of bile and blood throughout all experiments. After an initial 15-minute washout period, basal bicarbonate secretion (with luminal saline perfusion) was measured for 30 minutes. Subsequently, linaclotide (10^-9^ M, 10^-7^ M, 10^-5^ M) ± DRA_inh_-A250 inhibitor (10^-5^ M), CFTR_inh_-172 (2 × 10^-5^ M), or S3226 (10^-5^ M) was perfused intraluminally for 30 minutes. When multiple doses of drugs were studied, they were perfused sequentially in the same animals. These doses were selected based on previously published data and similarities with clinical dosing. Forskolin (10^-4^ M) ± CFTR_inh_-172 (2 × 10^-5^ M) was perfused intraluminally for 45 minutes. After each experiment, the length of the duodenal test segment was measured in situ to the nearest 0.5 mm.

Sample volumes were measured by weight to the nearest 0.01 mg. The amount of bicarbonate in the effluents was quantitated by a validated micro back-titration method.(36) Briefly, 100 μL of 5 × 10^-2^ M HCl was added to 2 mL sample with 2 mL of double-distilled water. Samples were then gassed with N_2_, prewashed in Ba(OH)_2_ to remove all CO_2_, and back-titrated with 2.5 × 10^-2^ M NaOH to an endpoint of pH 7.0 using an automated pH meter/titration unit (TIM 856 Radiometer, Copenhagen, Denmark). Bicarbonate secretion was determined in 15-minute periods and expressed as micromoles per centimeter per hour and presented as bicarbonate output over time or net peak bicarbonate output (peak output minus average basal).

### In Vitro Measurement of Ion Transport and Bicarbonate Secretion

For murine measurements, mice were anesthetized as above, and the proximal 1 cm of duodenum was removed and placed in ice-cold 3 × 10^-1^ M mannitol solution with indomethacin (10^-5^ M) to prevent endogenous prostaglandin formation. The mesentery was dissected off and the duodenum was opened along the mesenteric edge. The muscular layer was scraped from the mucosa using a glass slide while being bathed in the indomethacin-containing solution. Stripped duodenal mucosae were mounted between two 0.1 cm^2^ aperture sliders placed in temperature controlled Ussing chambers maintained at 37°C (P2300, Physiologic Instruments, San Diego, CA). For human endoscopic biopsy measurements, biopsies were collected in the same iso-osmolar mannitol solution with indomethacin and then mounted directly between two 0.031 cm^2^ aperture sliders. Tissue was bathed in apical solutions containing (in mM): NaCl 115, KGluconate 5.2, NaGluconate 25, MgCl_2_ 1.2, CaCl_2_ 1.2, Mannitol 10, gassed with 100% O_2_ and basolateral solutions containing (in mM): NaCl 115, K_2_HPO_4_ 2.4, KH_2_PO_4_ 0.4, NaHCO_3_ 25, MgCl_2_ 1.2, CaCl_2_ 1.2, Glucose 10, gassed with 95% O_2_/5% CO_2_. Ion transport was measured by short-circuit current (I_sc_), where tissues were voltage clamped at 0 mV using a voltage-clamp apparatus (VCC600, Physiologic Instruments, San Diego, CA). To monitor transepithelial resistance, the voltage clamp was released, and the open circuit voltage was recorded every 10 minutes. Resistance was calculated using Ohm’s Law. To measure bicarbonate secretion, we utilized the automatic pH-stat method, as previously described.(7, 37) In brief, a pH microelectrode was placed in the apical bath and set at the pH of the unbuffered apical solution (typically ∼6.8). The titrator (Titrando 902, MetrOhm, Switzerland) was set to titrate 5 × 10^-3^ M HCl in 0.2 μL aliquots as the pH of the apical bath rose above the pH set point. Tiamo software (MetrOhm, Switzerland) was used to control the rate of titration and continuously measure the amount of HCl titrated into the apical bath and the apical bath pH. The set-up as described allows for simultaneous measurement of bicarbonate secretion and I_sc_ without electrical interference. Bicarbonate secretory rates (μmol/cm^2^•h) were calculated in 5-minute intervals by noting the amount titrated, the concentration of titrant, and the surface area of the slider aperture. In a subset of experiments, forskolin (10^-4^ M, serosal) was added at the end of the experiment to assess tissue viability.

### Measurement of Intracellular pH

pH_i_ levels were measured via epiflorescent microscopy on a Leica Thunder DMi8 imaging system. Enteroids were grown on collagen-coated 18 mm coverslips in WENR (undifferentiated) alone or changed to EN for 3 days (differentiated) after initial seeding in WENR. On day of experiment, enteroids were incubated with the cell-permeant ratiometric pH indicator SNARF-5F 5-(and-6)-carboxylic acid, acetoxymethyl ester, acetate (5 µM) (Thermo-Fisher, Waltham, MA) for 30 minutes at room temperature. Following a 15-minute wash period, imaging experiments were performed in physiologic buffer (in mM: NaCl 115, K_2_HPO_4_ 2.4, KH_2_PO_4_ 0.4, MgCl_2_ 1.2, CaCl_2_ 1.2, HEPES 10, Glucose 10). Coverslips were imaged in a heat-controlled (37°C), gassed (95% O_2_/5% CO_2_) enclosure. Using LasX software, regions of interest (ROI) were selected by encircling visualized cells for ratiometric measurements within each ROI. Measurements were taken every 5 seconds. Baseline pH_i_ was recorded for 2-3 minutes, followed by application of CFTR_inh_-172 (20 × 10^-5^ M) or DRA_inh_-A250 (10^-5^ M, 5 min) then CFTR_inh_-172. Upon full pH_i_ recovery (typically 4-6 minutes), pH calibration was performed using serial addition of nigericin + valinomycin in pH 4.5 – 7.5 buffers (Thermo-Fisher, Waltham, MA) for at least 5 minutes each.

### Single cell sequencing analysis

The R toolkit Seurat was employed to conduct an analysis and quality control of single-cell RNA sequencing (scRNA-seq) data from existing literature.(38) We clustered the cells according to the Seurat default settings. In brief, we embedded the cells through a K-nearest neighbor graph, reduced the data with principal component analysis, and applied the Louvain algorithm for clustering. We visualized these clusters with UMAP (uniform manifold approximation and projection). To annotate major cell types and compare gene expression in crypts versus villi, we used the metadata annotations provided by Elmentaite et al.(21) and Busslinger et al.,(20) respectively. We utilized violin and feature plots to display gene expression profiles, and FeatureScatter to compare the co-expression of two genes by cell type.

### Human duodenal organoids

Duodenal organoids were established and cultured as previously described.(39) In short, crypt cells were isolated from duodenal endoscopic biopsies using a series of PBS washes followed by incubation with cold chelation buffer (distilled water with 5.6 mM Na_2_HPO_4_, 8.0 mM KH_2_PO_4_, 96.2 mM NaCl, 1.6 mM KCl, 43.4 mM sucrose, 54.9 mM d-sorbitol, 1 mM dl-dithiothreitol) with 2 μM EDTA for 30 mins on ice. After incubation, EDTA was removed, and the biopsies were vigorously resuspended in cold chelation buffer (without EDTA). The resuspension was allowed to sit on ice for 1 minute to allow biopsies to settle by gravity. The supernatant was collected. This process was repeated 3-5 times until no more crypts were present in the supernatant. The crypt containing supernatants were pooled and centrifuged at 600g for 5 minutes, followed by resuspension in 40 µL basement membrane extract /dome (BME, Culterex Pathclear Type 2 BME) and submerged in WENR complete media (ADMEM/F12, 1 mM HEPES, 1X Glutamax, 1 mM N-Acetylcysteine, 1X B-27, 0.5 µM A83-01, 1X Penicillin-Streptomycin-Glutamine, 10 mM Gastrin, 10 µM SB-202190, 50 ng/mL EGF, 10 mM Nicotinamide, 1X Normocin) with Rho-associated kinase inhibitor (Y-27632 10 µM, Sigma Aldrich, St. Louis, MO) and CHIR99021 (10µM, Sigma Aldrich, St. Louis, MO) for first two days, followed by WENR complete media only until ready to passage. Duodenal enteroids were passaged by disrupting BME with cold (4°C) media, 1X cold (4°C) phosphate-buffer solution (PBS, without Ca^2+^ and Mg^2+^), then centrifuged at 600g for 3 minutes at 4°C. Supernatant was aspirated and single cells were isolated by adding warm (37°C) TrypLE Express (Gibco, Thermofisher Scientific, Waltham, MA) and incubated for 10 minutes at 37°C. Equal volume of fetal bovine serum was added to quench trypsin and cells were centrifuged at room temperature for 5 minutes at 600g. Pellet was resolubilized in BME (40 μL/dome) and plated in a 24-well tissue culture plate. Generation of apical-out enteroids was performed as previously described.(22) Briefly, enteroids were dislodged from BME using with 1X cold (4°C) PBS (without Ca^2+^ and Mg^2+^) and resuspended in 5 mM EDTA in PBS for 1 hour at 4°C. Enteroids were then centrifuged at 200g for 3 minutes and resuspended in WENR or EN media (WENR without Wnt and R-Spondin) and seeded in ultra-low-attachment 24-well plates (Corning Costar, Thermofisher Scientific, Waltham, MA) for three days. On Day 3, undifferentiated and/or differentiated enteroids were harvested for experiments.

### Quantitative PCR

Quantification of mRNA transcript levels in undifferentiated and differentiated apical-out enteroids was performed. Briefly, total RNA was extracted from fresh apical-out enteroids using Zymo Direct-Zol RNA miniprep kit (Zymo Research, Irvine, CA) according to the manufacturer’s instructions. Complementary DNA (cDNA) was synthesized from 1 to 2 μg of RNA using High-Capacity RNA-to-cDNA™ Kit (Zymo Research, Irvine, CA). Quantitative real-time polymerase chain reaction (qPCR) was performed on a StepOne Plus real-time PCR system (Applied Biosystems, Foster City, CA) using PowerUP SYBR Green Master Mix (Applied Biosystems, Foster City, CA). Each sample was run in triplicate in a 20 μL reaction volume containing 5 ng cDNA/reaction. PCR was carried out for 40 cycles at 95°C (denature) for 30 seconds, 60°C (anneal/extend) for 30 seconds and amplification data was collected after each cycle. The sequences of optimized gene-specific primers are listed in Supplemental Table 1. Glyceraldehyde-3-phosphate dehydrogenase (*GAPDH*) was used as the reference gene to obtain delta threshold cycles (ΔCt) values. The relative expression levels of the target genes were calculated using the 2^-ΔΔCt^ method.

### Confocal microscopy

Treated Day 3 apical-out enteroids were fixed in 2% paraformaldehyde immediately after treatment and stored at 4°C until processing. All steps were performed in ultra-low attachment plates. Fixed enteroids were permeabilized for 30 minutes with permeabilization buffer (3% bovine serum albumin, BSA, ThermoFisher Scientific, Waltham, MA), 0.5% Triton X-100, and 0.7% Saponin (Sigma-Aldrich, St. Louis, MO) in PBS and blocked for 60 minutes with gentle shaking in blocking buffer (3% BSA, 0.2% Triton X-100). Samples were incubated overnight at 4°C with primary antibodies (Supplemental Table 2) diluted in antibody diluent (3% BSA, 0.2% Triton X-100, and 0.5% Saponin in PBS), washed 4 times, 30 minutes each in antibody diluent buffer, and incubated with secondary antibodies (Table 2) for 1 hour with gentle shaking at room temperature. Samples were washed 4 times for 30 minutes each in antibody diluent buffer. Samples were then carefully resuspended in SlowFade gold antifade mounting medium (ThermoFisher Scientific, Waltham, MA), mounted on microscope slides and sealed with glass coverslips (1.0 mm) (ThermoFisher Scientific, Waltham, MA). Slides were imaged using Leica Stellaris 8 Confocal microscope. Z-stacks of 0.33 μm sections/sample were obtained and analyzed using FIJI Software (ImageJ). Following acquisition, image settings were adjusted to optimize the signal to noise ratio. For quantitative analysis, the same settings were used across the different conditions compared, and the protein-of-interest mean fluorescence intensity (MFI) after background subtractions was analyzed. Villin was used as a marker for the plasma membrane and used to identify the area for fluorescence intensity measurements. Drug-treated MFI measurements were normalized to corresponding vehicle control MFI measurements.

### Statistics

Results are expressed as means ± standard error of the mean (SEM) for a series of n experiments. Statistical analysis was performed by using the Student’s t-test for unpaired data or by ANOVA with repeated measures analysis, as appropriate. P values of <0.05 were considered significant.

### Study Approval

All animal protocols were approved by the Institutional Animal Care and Use Committee at Stanford University (#33183). Children and adults undergoing upper endoscopy for clinical indications were approached to obtain additional duodenal biopsies for research under an Institutional Review Board approved protocol (#47536). Subjects without endoscopic and microscopic evidence of duodenal pathology were included in this study.

## AUTHOR CONTRIBUTIONS

JBS: Performed and analyzed experiments, drafted, edited and approved final manuscript.

AMT: Performed and analyzed experiments, drafted and approved final manuscript.

SMA: Performed and analyzed experiments, drafted and approved final manuscript.

VVU: Performed and analyzed experiments, drafted and approved final manuscript.

JEC: Performed and analyzed experiments, approved final manuscript.

YF: Performed and analyzed experiments, approved final manuscript.

JG: Performed and analyzed experiments, approved final manuscript.

MOA: Contributed technical expertise, material support, approved final manuscript.

OC: Contributed technical expertise, material support, approved final manuscript.

CJK: Contributed technical expertise, material support, edited and approved final manuscript.

ZMS: Study conceptualization, experimental design, funding, drafted, edited and approved final manuscript.

## ACKNOWLEDGEMENTS

Drs. Anjaparavanda Naren and Kyu Shik Mun (Cedars Sinai) kindly provided CF enteroids. We thank Dr. Manuel Amieva (Stanford University) for technical advice on apical-out enteroids. This research was supported by the National Institute of Diabetes, Digestive, and Kidney Diseases (K08DK124684 to ZMS, U01DK085527 and R01DK115728 to CJK), R01DK126070 and P30DK072517 to OC, the Cystic Fibrosis Foundation (SELLER19GE0, SELLER20-KB, SELLER16L0 to ZMS and CIL21G0 to OC), North American Society of Pediatric Gastroenterology, Hepatology, and Nutrition Foundation (ZMS), Stanford University (ZMS), Stanford Maternal Child Health Research Institute (ZMS), and Sellers Research and Clinical Development, LLC (ZMS).

## DATA AVAILABILITY

The authors declare that the data supporting the findings of this study are available within the paper and its supplementary information files. Any further information on the data in this study are available by reasonable request to the corresponding author.

## PREPRINT

This manuscript has been deposited on a public preprint server (doi: https://doi.org/10.1101/2023.05.05.539132).

