## Supplementary Tables and Figures for "Key role of down-regulated in adenoma (*SLC26A3*) chloride/bicarbonate exchanger in linaclotide-stimulated intestinal bicarbonate secretion upon loss of CFTR function"

**Supplementary Table 1. qPCR gene-specific Primer sequences**

| Gene | Forward Primer (5′-3′) | Reverse Primer (5′-3′) |
| --- | --- | --- |
| *SLC26A3* | CAGCCCCCTATTACACCTGA | CCTCCTGTGCTCTCCTGAAC |
| *SLC26A6* | GCCTTGAACGACTCCATGAT | TGTGAGACGAAGACCTGCAC |
| *CFTR* | CCTATGACCCGGATAACAAGGA | GAACACGGCTTGACAGCTTTA |
| *SLC9A3* | CCTGACCATCAAGCCTCTGG | ACATTCAGGATCCGGTCTCG |
| *LGR5* | GATGTTGCTCAGGGTGGACT | GGGAGCAGCTGACTGATGTT |
| *GUCY2C* | GGCTGTCCTTTAGTTCCCAGG | GAAAGTAGCGTTCACAGTCACAT |
| *MYO6* | TAACCCACTCCTAGAAGCCTTT | GCACCAGCACACAACCTATAA |
| *GAPDH* | GAAGGTGAAGGTCGGAGTC | GAAGATGGTGATGGGATTTC |
| SLC26A3 (down-regulated in adenoma, DRA), SLC26A6 (putative anion transporter-1, PAT-1), CFTR (cystic fibrosis transmembrane conductance regulator), SLC9A3 (sodium/hydrogen exchanger 3, NHE3), LGR5 (leucine rich repeat containing G protein-coupled receptor 5), GUCY2C (guanylate cyclase 2C), MYO6 (myosin VI), GAPDH (glyceraldehyde-3-phosphate dehydrogenase) | | |

**Supplementary Table 2. List of antibodies**

| Primary Antibodies | Dilution | Manufacturer |
| --- | --- | --- |
| DRA (mouse) | 1:200 | Santa Cruz (sc-376187) |
| NHE3 (rabbit) | 1:400 | Novus Biologicals (NBP1-82574) |
| Villin (rabbit) | 1:400 | ThermoFisher (PA5-29078) |
| Villin (mouse) 1D2C3 | 1:200 | Santa Cruz (sc-58897) |
| Hoechst 33342 | Per manufacturer’s protocol | Abcam (ab228551) |

| Secondary antibodies | Dilution | Manufacturer |
| --- | --- | --- |
| Goat Anti-Rabbit Alexa flour 594 | 1:300 | Abcam (ab150088) |
| Goat Anti-Mouse Alexa flour 647 | 1:300 | Abcam (ab150115) |
| Goat Anti-Mouse IgG H&L (Alexa Fluor® 488) | 1:1000 | Abcam (ab150113) |

**SUPPLEMENTARY FIGURES**

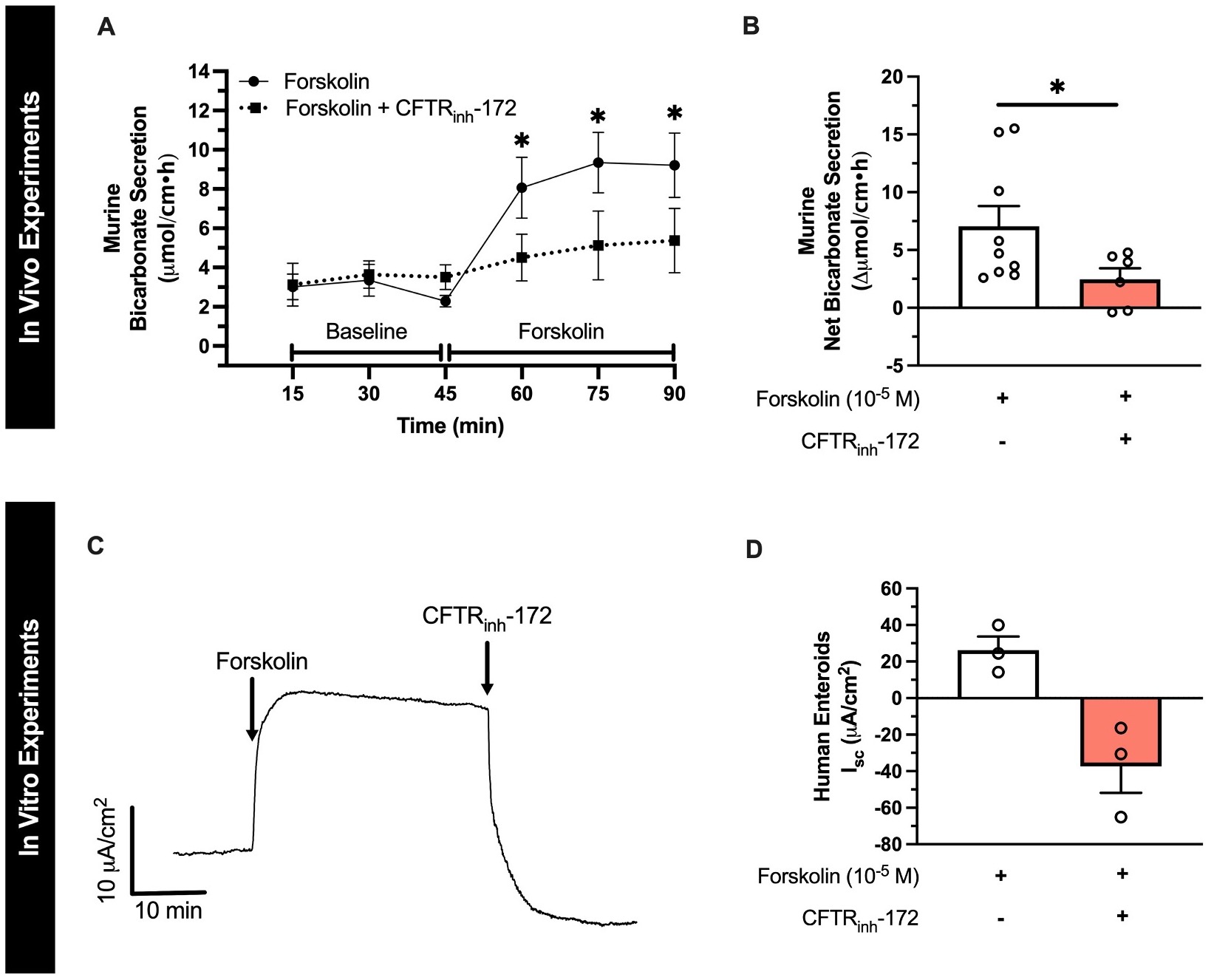

**Supplemental Figure 1. CFTR_inh_-172 inhibits forskolin-stimulated duodenal bicarbonate secretion in mice and human enteroids.** **A.** To compare the effect of pharmacologic CFTR inhibition with CFTR_inh_-172 on linaclotide and forskolin-stimulated bicarbonate secretion, we performed similar in vivo experiments as Figure 2A. Duodenum of wildtype mice were perfused with saline then forskolin (10^-4^ M) (black circles) or saline + CFTR_inh_-172 (2 x 10^-5^ M) then forskolin (10^-4^ M) (black squares). Bicarbonate secretion was calculated from the perfusates similar to Figure 2A. *, P<0.05 by ANOVA. **B.** Comparison of net change in forskolin-stimulated bicarbonate secretion (peak response – baseline secretion) in the presence or absence of CFTR_inh_-172. Columns with whiskers represent mean ± SEM with each circle representing a different experiment. Mean ± SEM percent inhibition indicated above columns. *, P<0.05 by unpaired Student’s t-test. **C.** Representative trace of forskolin (10^-5^ M, basolateral)-stimulated short-circuit current, followed by inhibition by CFTR_inh_-172 (2 x 10^-5^ M, basolateral), in undifferentiated human duodenal enteroids monolayers. D. Quantification of replicate experiments performed in C. Columns with whiskers represent mean ± SEM with each circle representing a different experiment.

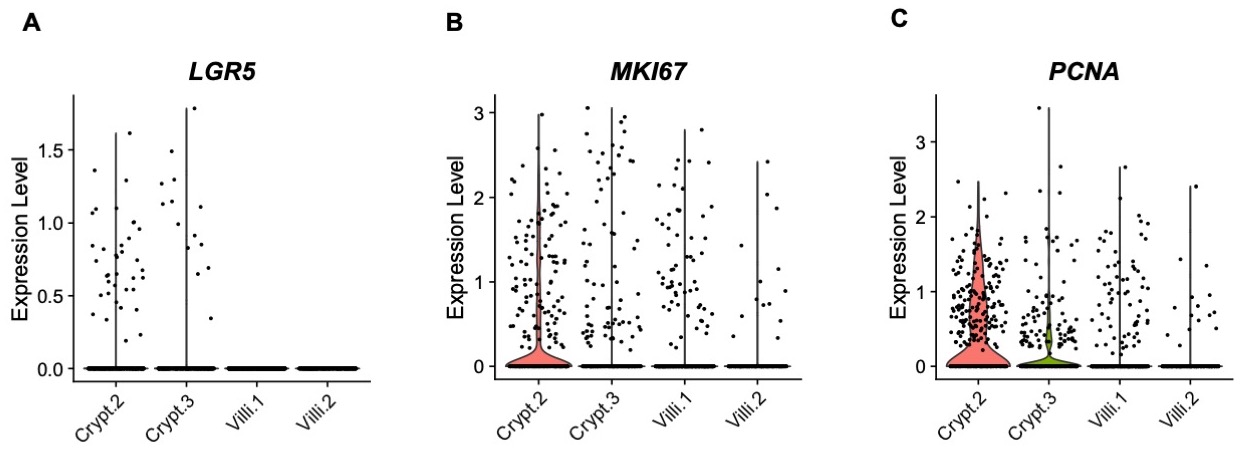

**Supplementary Figure 2. Validation of previously published human duodenum scRNA-seq datasets.** **A-C.** To verify the crypt vs. villi identity in the Busslinger et al. dataset we examined the expression of *LGR5* **(A)**, *MKI67* **(B)**, and *PCNA* **(C)**.

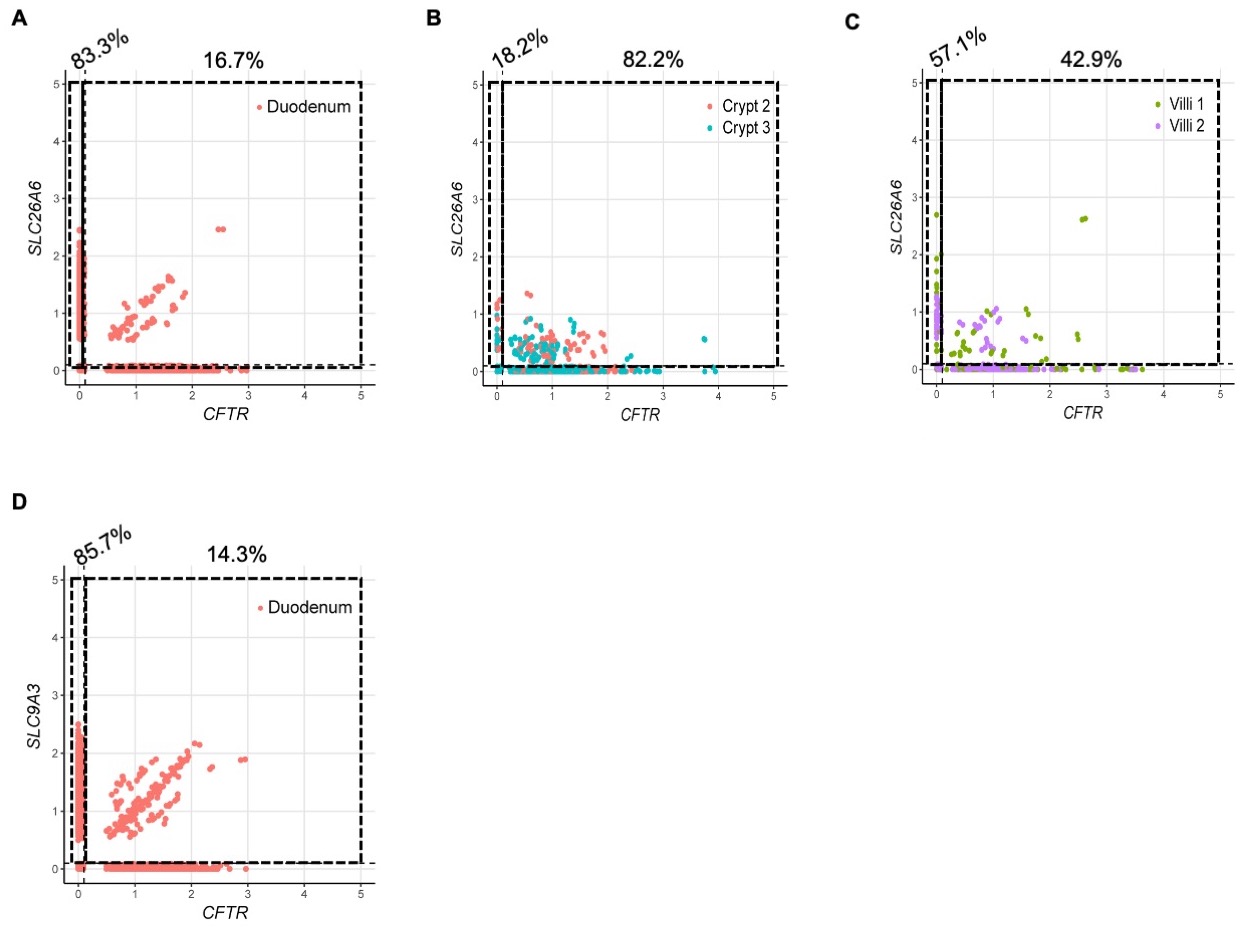

**Supplementary Figure 3. Co-expression of *CFTR* and *SLC26A6* (PAT-1) and *SLC9A3* (NHE3) from human duodenal sc-RNAseq data.** **A-C.** Co-expression of *SLC26A6* (PAT-1) and *CFTR* mRNA using FeatureScatter based on Elmentaite et al.^14^ enterocytes (A) and Busslinger et al.^15^ crypt and villi (B and C) datasets. **D.** Co-expression of *SLC9A3* (NHE3) and *CFTR* mRNA using FeatureScatter based on Elmentaite et al. enterocytes. There were insufficient numbers of *SLC9A3* positive cells in the Busslinger et al. dataset to do the same analysis as B and C. Numbers on top of graphs represent the percentage of *SLC26A6*- or *SLC9A3*-expressing cells that express these only (left) or *SLC26A6/SLC9A3* and *CFTR* (right).

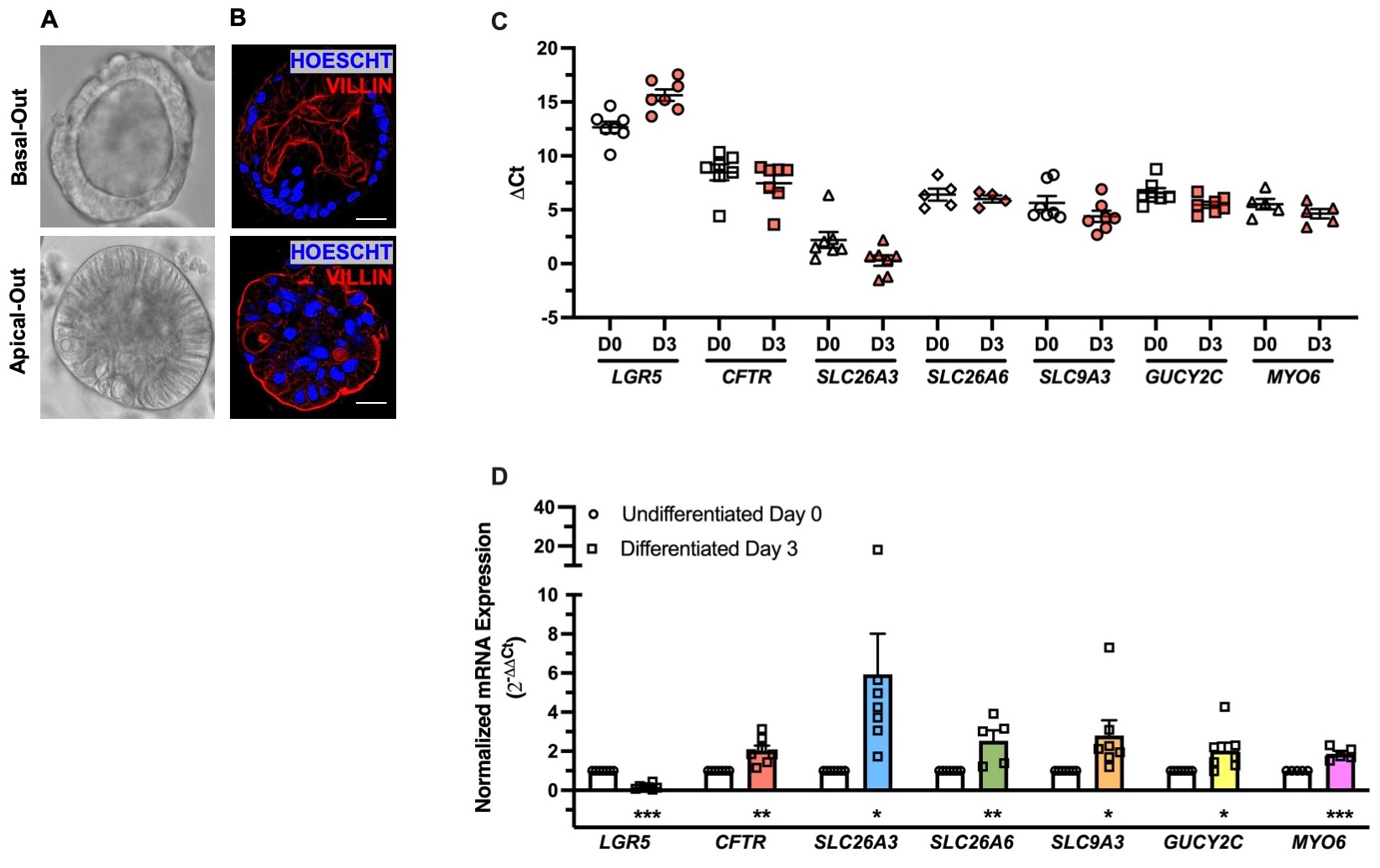

**Supplementary Figure 4. Characterization of Apical-Out Duodenal Enteroids. A.** Sample brightfield microscope images of basal-out (top) and apical-out (bottom) human duodenal enteroids. **B.** Representative confocal immunofluorescent images of basal-out (top) and apical-out (bottom) human duodenal enteroids stained with villin (apical membrane) and Hoescht (cell nucleus). Scale bar = 20 μm. **C and D.** mRNA expression of *LGR5, CFTR, SLC26A3, SLC26A6, SLC9A3, GUCY2C, and MYO6* in undifferentiated and differentiated (x3 days) apical-out human duodenal enteroids. Delta CT was calculated by comparing to *GAPDH* expression (C) and Delta Delta CT was calculated by comparing Differentiated Day 3 enteroids to Undifferentiated Day 0 enteroids. *, P<0.05; **, P<0.01; ***, P<0.001 compared to Undifferentiated Day 0 enteroids by unpaired Student’s t-test.

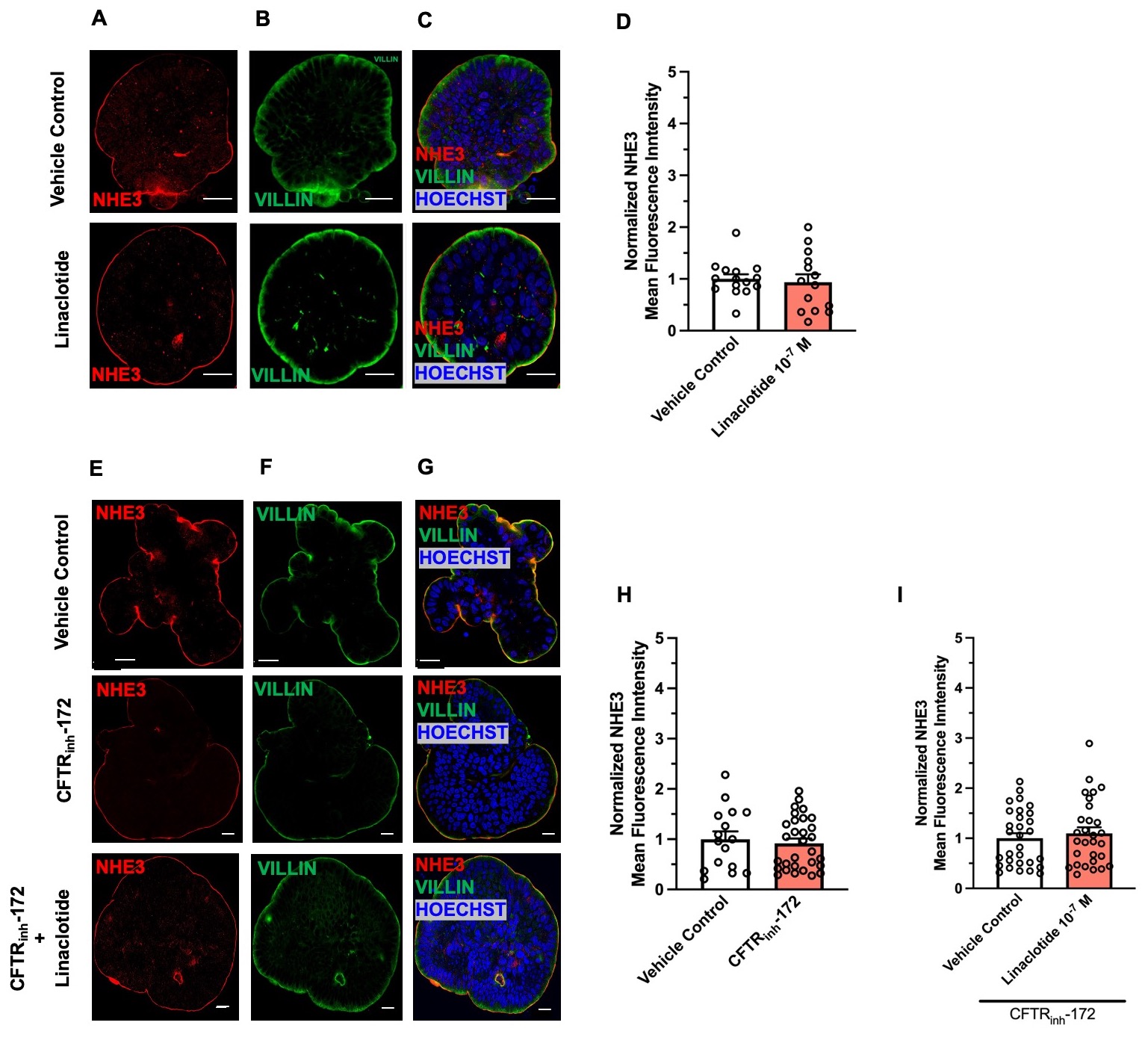

**Supplemental Figure 5. Linaclotide does not change the membrane expression of NHE3 in apical-out human duodenal enteroids. A-C.** Representative confocal microscopy images of NHE3 (A), villin (B), and NHE3 and villin and Hoescht (C) during control conditions (water, 40 minutes, top) or linaclotide (10^-7^ M, 40 minutes, bottom). **D.** Quantification of NHE3 present at the apical membrane using villin to define apical membrane, similar to Fig. 6. **E-G.** Representative images for apical membrane NHE3 mean fluorescence intensity following CFTR_inh_-172 (2 x 10^-5^ M, 40 minutes) with or without linaclotide (10^-7^ M, 40 minutes) treatment. H-I. Quantification of images, normalized to vehicle controls for comparison. Vehicle control for linaclotide was water and for CFTR_inh_-172 was DMSO. Columns with whiskers are mean ± SEM with each dot representing a different enteroid. Enteroids from three different patients were used for each condition. Significance determined by unpaired Student’s t-test. For enteroids treated with both CFTR_inh_-172 and linaclotide, enteroids were pretreated with CFTR_inh_-172 for 40 minutes prior to linaclotide treatment.
